# Selective autophagic clearance of protein aggregates is mediated by the autophagy receptor, TAX1BP1

**DOI:** 10.1101/558767

**Authors:** Shireen A. Sarraf, Hetal V. Shah, Gil Kanfer, Michael E. Ward, Richard J. Youle

## Abstract

Misfolded protein aggregates can disrupt cellular homeostasis and cause toxicity, a hallmark of numerous neurodegenerative diseases. Protein quality control by the ubiquitin proteasome system (UPS) and autophagy is vital for clearance of aggregates and maintenance of cellular homeostasis^1^. Autophagy receptor proteins bridge the interaction between ubiquitinated proteins and the autophagy machinery allowing selective elimination of cargo^2^. Aggrephagy is critical to protein quality control, but how aggregates are recognized and targeted for degradation is not well understood. Here we examine the requirements for 5 autophagy receptor proteins: OPTN, NBR1, p62, NDP52, and TAX1BP1 in proteotoxic stress-induced aggregate clearance. Endogenous TAX1BP1 is both recruited to and required for the clearance of stress-induced aggregates while overexpression of TAX1BP1 increases aggregate clearance through autophagy. Furthermore, TAX1BP1 depletion sensitizes cells to proteotoxic stress and Huntington’s disease-linked polyQ proteins, whereas TAX1BP1 overexpression clears cells of polyQ protein aggregates by autophagy. We propose a broad role for TAX1BP1 in the clearance of cytotoxic proteins, thus identifying a new mode of clearance of protein inclusions.

---

Maintenance of cellular and organismal health is intricately connected to protein quality control. A balance exists between protein translation, folding, and degradation that maintains the stoichiometry and function of cellular protein complexes and organelles. If this balance is perturbed, the accumulation of misfolded proteins can be toxic to the cell and is associated with disruption of cellular function and numerous neurodegenerative diseases such as amyotrophic lateral sclerosis (ALS), Huntington's disease (HD), and Alzheimer’s disease (AD)^3^. A number of protein quality control pathways exist within the cell to forestall dysfunction. Molecular chaperone systems function to monitor and refold proteins if possible, while misfolded or damaged proteins are targeted for elimination via autophagy or the UPS^4,5^. The UPS is generally responsible for routine turnover of short-lived proteins and targeted degradation of soluble or solubilized misfolded proteins while macroautophagy, a catabolic process, culminates in the lysosomal degradation of long-lived proteins, damaged organelles, and portions of the cytoplasm^6–9^. Since the proteasome can only accommodate unfolded polypeptide chains, it is generally thought that autophagy is responsible for the removal of insoluble protein aggregates^1,10^. If either autophagy or the UPS is hindered, acute or chronic proteotoxic stress, such as that caused by the expression of mutated proteins in neurodegenerative disease, can result in the selective accumulation of these aggregation-prone proteins^11–13^. *In vivo*, inhibition of autophagy results in intracellular protein aggregation contributing to neuronal cell death and neurodegeneration in mice^14,15^.

In recent years, numerous studies have highlighted the ability of autophagy to selectively eliminate specific substrates, including mature ribosomes, endoplasmic reticulum, intracellular pathogens, mitochondria, and protein aggregates^16–19^. In these cases, the autophagic machinery employed in nonselective bulk degradation of cytosolic material is targeted to specific cargo. Stimulation of autophagy is a promising therapeutic strategy in the treatment of protein aggregation diseases and has been shown to enhance turnover of aggregated proteins, such as TDP-43, in neuronal ALS models and huntingtin protein in Huntington disease models^20–22^. Key to this approach is an understanding of how autophagy is selectively targeted to aggregates. In addition to the specificity mediated by E3 ubiquitin (UB) ligases that target their cognate substrates, there is selectivity in delivery to the proteasome via UB receptors^23^ and in the recruitment of autophagic machinery via autophagy receptors which associate with polyubiquitylated proteins, thus linking substrates to the appropriate degradation machinery^16^.

Numerous types of selective autophagy have been examined, and much progress has been made in determining the basis of cargo selectivity. The autophagy receptor proteins, OPTN, NDP52, TAX1BP1 and p62 are known to be important in xenophagy^24,25^, while OPTN, NDP52, and to a lesser extent, TAX1BP1 are essential for PINK1/Parkin-mediated mitophagy^17,26,27^. A screen in yeast identified the ATG8 and UB-binding protein, CUET, and its mammalian homologue, TOLLIP, to contribute to autophagic removal of expanded polyQ isoforms of huntingtin^28^. NBR1 and p62 are linked to aggrephagy in mammalian cells^29^; however, p62 and NBR1 knockout (KO) mice exhibit only a mild increase in ubiquitylated aggregates suggesting that additional receptors are involved^30^. To identify such autophagy receptor proteins, we examined production and clearance of puromycin-induced truncated and misfolded proteins in the cytosol^31^, termed “DRiPs” for **d**efective **ri**bosomal **p**roducts^32^ in individual and combinatorial knockouts of the autophagy receptor proteins, NBR1, p62, NDP52, OPTN, and TAX1BP1 (Fig. 1A, 1B, 1C, Supplementary Fig. 1A). Consistent with the literature^33,34^, we found a drastic reduction in puromycin-induced foci formation in p62 KO and NBR1 KO cell lines, as well as in NDP52 KO cells compared to WT cells after 2h puromycin (Fig. 1C, 1D, 1E). Though aggregate formation was decreased in NBR1 and NDP52 KO cell lines, any foci that did form were effectively cleared after 3h washout of puromycin (Fig. 1C, 1F, 1G). In contrast, individual OPTN KO and TAX1BP1 KO cell lines showed robust UB-positive foci formation, equal to or greater than WT cells after 2h puromycin (1C, 1D, 1E) and a significant block in aggregate clearance after puromycin washout. More than twice as many cells with foci (~54%) persisted in OPTN KO cells compared to WT (~22%), while in TAX1BP1 KO cells, nearly 4 times as many cells retained foci (~85%) (Fig. 1C, 1F, 1G). In TAX1BP1 KO cells with foci, the number of foci per cell, although equivalent to WT at 2h puromycin (Supplementary Fig. 1B, 1C), was substantially higher than WT cells upon puromycin washout, indicating a block in clearance (Supplementary Fig. 1D). Distinct roles therefore exist for autophagy receptors in protein aggregate formation and clearance and our data reveal a new autophagy receptor involved in aggregate clearance.

**Figure 1.**
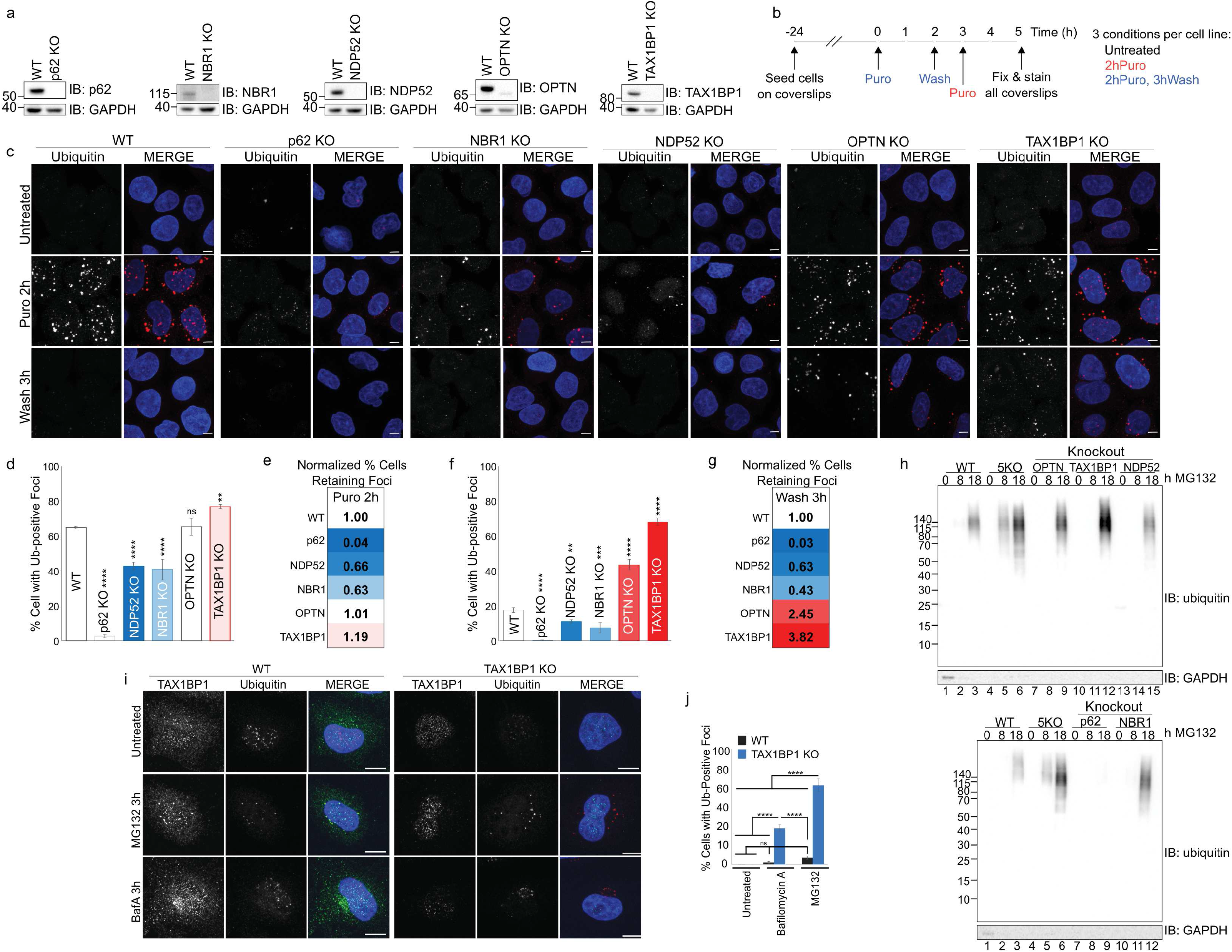
TAX1BP1 depletion impairs clearance of protein aggregates. **a,** Validation of knockout cell lines. **b,** Experimental outline for assessing aggregate formation and clearance. **c,** WT or individual knockouts for p62, NBR1, NDP52, OPTN, and TAX1BP1 cell lines were exposed to 5 μg/ml puromycin for 2 h, after which cells were either fixed for imaging or washed and followed for a further 3 h in full media; scale bar 10μm. **d, f,** Quantification of b: percent of cells containing UB-positive foci was assessed in ~ 200 cells per condition in 3 independent experiments at (d) 2 h puromycin or (f) 2 h puromycin followed by 3 h washout. Quantification is displayed as mean ± s.d. from 3 independent experiments using one-way ANOVA test (\*\**P*<0.01, \*\*\**P*<0.001,\*\*\*\**P*<0.0001) comparing all to WT and Tukey’s post hoc test. **e, g,** WT normalized comparisons of foci formation and foci clearance in autophagy receptor knockout cell lines. h, WT, pentaKO, or individual KO lines for each autophagy receptor were treated with 1 μM MG132 for 8 or 18 **h,** fractionated into RIPA-soluble or -insoluble fractions and immunoblotted for total ubiquitin. Quantification and soluble fractions shown in Supp. Fig. 1C and 1D, respectively. **i,** WT or TAX1BP1 KO cell lines were exposed to 100 nM Bafilomycin A or 1 μM MG132 for 5 h then fixed for imaging. **j,** Quantification of i: percent of cells containing UB-positive foci was assessed in ~ 200 cells per condition. Quantification is displayed as mean ± s.d. from 3 independent experiments using one-way ANOVA test, (\*\*\*\**P*<0.0001) comparing all to WT and Tukey’s post hoc test. The number of observations used in each experimental series and *P* values for all comparisons are included in Table 1. All blots and microscopy images are representative of at least 3 independent experiments.

We observed complementary results upon exposure of cells to low levels of proteasome inhibition which induces accumulation of misfolded protein into juxtanuclear aggresome-like structures removed via autophagy^1^. Imaging revealed decreased UB-labeled foci formation in p62 KO, NBR1 KO and NDP52 KO cells (Supplementary Fig. 1E). Similarly, immunoblot of cellular lysates after fractionation revealed no accumulation of insoluble UB-positive protein in the p62 KO and decreased levels in NBR1 KO and NDP52 KO cells relative to WT cells (Fig. 1H, Supplementary Fig. 1F, 1G). However, in OPTN KO and TAX1BP1 KO cells, imaging revealed greater UB-foci accumulation than in WT cells (Supplementary Fig. 1E). Consistent with these observations, UB-conjugated protein was increased in the insoluble fraction of TAX1BP1 KO (~4X increase) and OPTN KO (~2X increase) compared to WT cells (Fig. 1H, Supplementary Fig. 1F), consistent with a block in aggrephagy.

We compared accumulation of UB-positive punctae upon short term exposure to MG132 or Bafilomycin A, an autophagy inhibitor, to determine whether there is basal accumulation of misfolded or aggregated protein in TAX1BP1 KO cells. We saw little effect in WT cells (Fig. 1I, 1J); but in TAX1BP1 KO cells, both inhibitors resulted in formation of UB-positive foci (Fig. 1I, 1J), though MG132 to a much greater extent (Fig. 1I, 1J). This implies that the UPS is compensating for an increased basal proteotoxic load in TAX1BP1 KO cells, most likely due to a block in TAX1BP1-mediated selective aggrephagy. Interestingly, proteotoxic stress induced a dramatic increase in TAX1BP1 protein levels (Fig. 2A, 2B). Additionally, a large proportion of TAX1BP1 accumulates with insoluble ubiquitinated protein after MG132 treatment (Fig. 2C).

**Figure 2.**
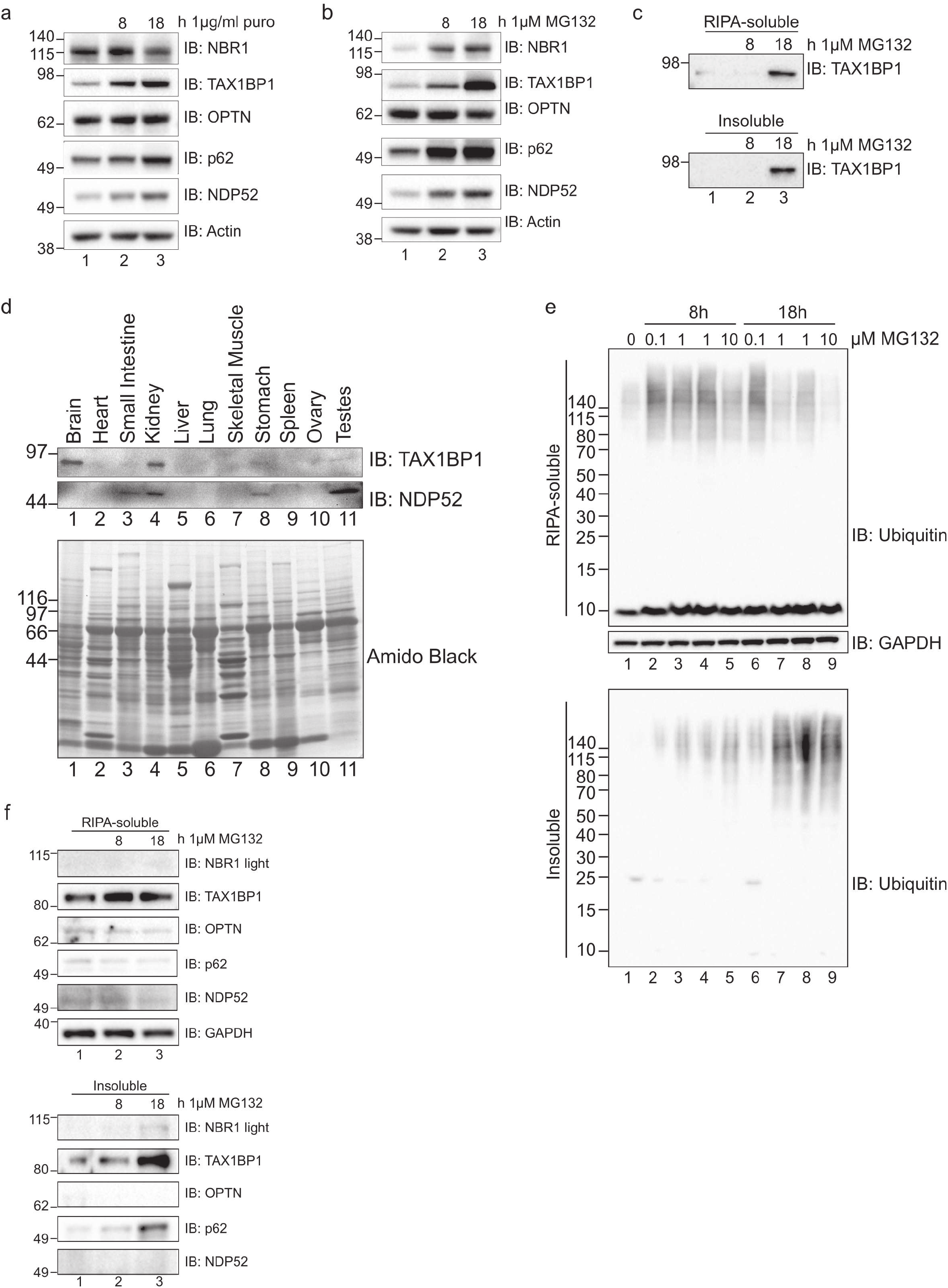
TAX1BP1 protein responds to proteotoxic stress and associates with insoluble protein. **a, b,** WT HeLa cells were treated with either (a) puromycin or (b) MG132 as indicated, lysed in 2% SDS and immunoblotted for TAX1BP1, OPTN, NBR1, p62, or NDP52. **c,** WT cells treated with 1 μM MG132 for 8 or 18 h were fractionated into RIPA-soluble or -insoluble fractions and immunoblotted for TAX1BP1. **d,** Human tissue panel probed for TAX1BP1 or NDP52. **e,** Primary rat cortical neurons were treated with the indicated amounts of MG132 for 8 or 18 h, fractionated into RIPA-soluble or -insoluble fractions and immunoblotted for total UB. **f,** Fractionated primary rat cortical neurons treated as in (e) were blotted for the indicated proteins. All blots are representative of at least 3 independent experiments.

TAX1BP1, NDP52, and CALCOCO1 are paralogous proteins of the CALCOCO gene family, all containing a SKICH domain, a putative canonical LC3-interacting region (LIR) motif and coiled-coil regions while differing in conservation of the C-terminal zinc finger domains in various species^25^. Therefore, we suggest that the specificity of TAX1BP1 and NDP52 may have diverged over time, leading to a specific function for TAX1BP1 in aggrephagy. TAX1BP1 is the most evolutionarily conserved member of the CALCOCO gene family and present widely in vertebrates, unlike NDP52 which is sporadically lost or truncated^25^. Examination of NDP52 and TAX1BP1 expression levels in a panel of human tissue lysates revealed that TAX1BP1 is highly expressed in brain, while NDP52 was undetectable and likely does not function in neuronal tissues (Fig. 2D). We then examined primary rat cortical neurons exposed to proteotoxic stress. As in HeLa cells, UB-conjugated protein accumulated in the insoluble fraction upon MG132 exposure (Fig. 2E). As in human tissue, TAX1BP1 was expressed robustly in the rat cortical neuron lysate and furthermore, accumulated to a greater extent in the insoluble fraction with increased exposure to proteotoxic stress compared to other autophagy receptor proteins except p62 (Fig. 2F). Consistent with our observations in HeLa cells, TAX1BP1 staining was diffuse in untreated primary rat cortical neurons and in neurons derived from human induced pluripotent stem cells (iPSCs)^35^, but colocalized robustly with UB upon MG132 treatment (Supplemental Fig. 2A, 2B). To our knowledge, this is the first study to identify a role for TAX1BP1 in aggrephagy and suggests that TAX1BP1 plays a specific role in neuronal aggrephagy.

To further validate a specific role for TAX1BP1 in aggrephagy, we reintroduced GFP-TAX1BP1 at near endogenous levels in the TAX1BP1 KO HeLa cell line (Supplementary Fig. 3A). In untreated cells, GFP-TAX1BP1 was diffuse or present in small punctae (Fig. 3A) and colocalized with UB-positive foci upon 2h puromycin treatment (Fig. 3A). After puromycin washout, UB foci were cleared to a similar extent in the TAX1BP1-rescued TAX1BP1 KO cells as in WT cells and GFP-TAX1BP1 returned to a diffuse or punctate appearance (Fig. 3A, 3B).

**Figure 3.**
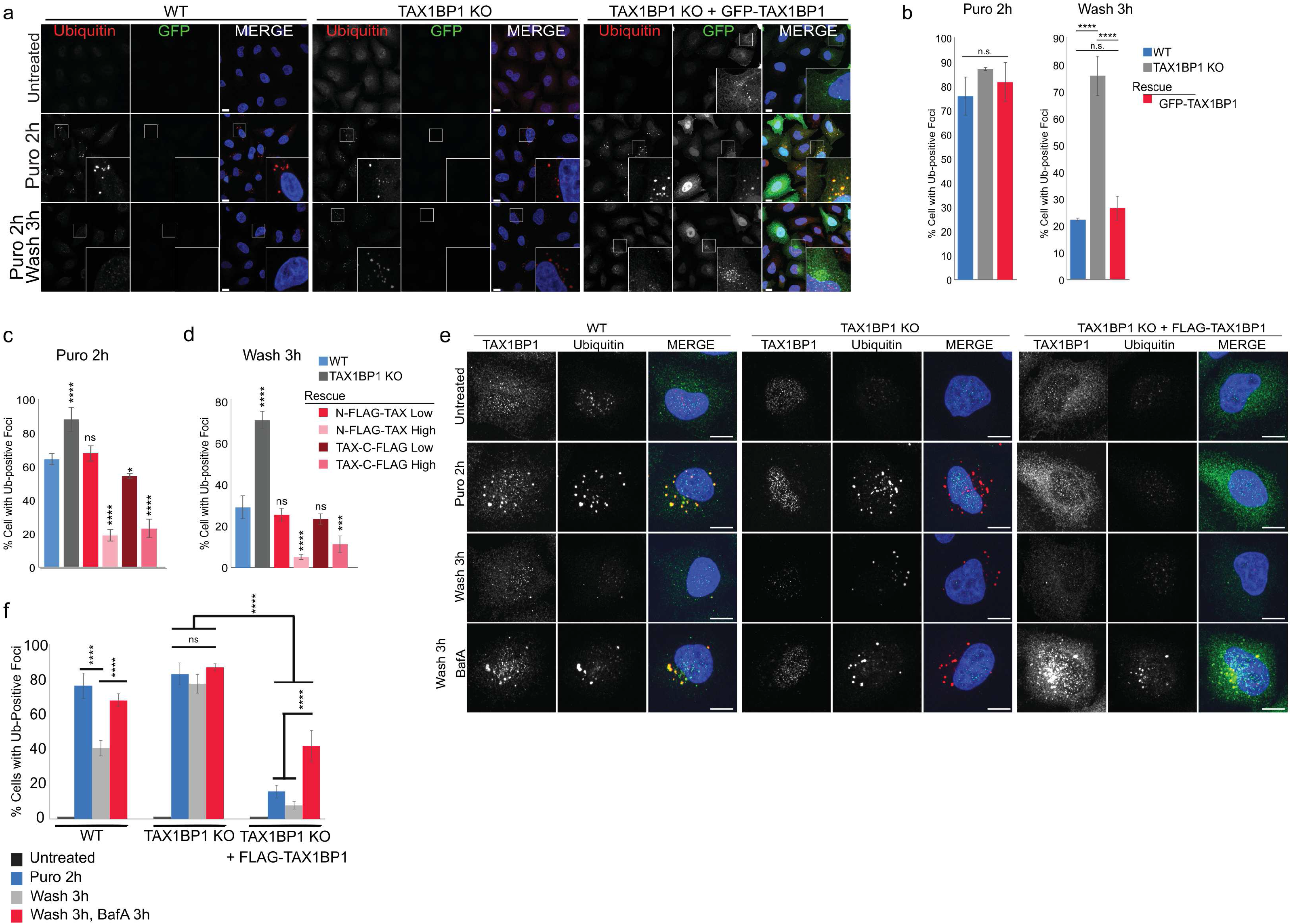
TAX1BP1 mediates aggregate clearance. **a,** WT, TAX1BP1 KO, and TAX1BP1 KO with stable expression of GFP-TAX1BP1 rescue cell lines were exposed to 5 μg/ml puromycin for 2 h, after which cells were either fixed for imaging or washed and followed for a further 3 h in full media; scale bar 10μm. **b,** Quantification of (a): percent of cells containing UB-positive foci was assessed in ~ 200 cells per condition. Quantification is displayed as mean ± s.d. from 3 independent experiments using one-way ANOVA test (\*\*\*\**P*<0.0001) and Tukey’s post hoc test. **c, d,** Stably-expressing TAX1BP1 rescue lines were created using either N-FLAG or C-FLAG tag at high (H) or low (L) expression levels (See Supplementary Figure 3A, 3B) and exposed to 5 μg/ml puromycin for 2 h, after which cells were either fixed for imaging (c) or washed and followed for a further 3 h in full media (d) and quantified as in (b). **e,** WT, TAX1BP1 KO, or TAX1BP1 KO + FLAG-T AX 1BP1 (H) cell lines were exposed to 5 μg/ml puromycin in the presence or absence of 100 nM Bafilomycin A, after which cells were either fixed for imaging or washed and followed for a further 3 h in full media or in full media containing Bafilomycin A. Larger fields of view shown in Supplementary Figure 3C. **f,** Quantification of e: percent of cells containing UB-positive foci was assessed in ~ 200 cells per condition in 3 independent experiments. Quantification is displayed as mean ± s.d. from 3 independent experiments using two-way ANOVA test, (\*\*\*\**P*<0.0001). The number of observations used in each experimental series and *P* values for all comparisons are included in Table 1. All images are representative of at least 3 independent experiments.

To further confirm the effect of TAX1BP1 on aggregate clearance, we titered expression of N- or C-terminally tagged TAX1BP1 constructs in the TAX1BP1 KO background to create stable high- or low-expressing rescue lines (Supplementary Fig. 3A, 3B). Aggregate formation and clearance was similar to WT in the low-expressing TAX1BP1 rescue lines (Fig. 3C, 3D, Supplementary Fig. 3B). However, upon 2h puromycin treatment, UB-positive foci were present in only ~20% of the high TAX1BP1-expressing rescue cell lines compared to ~65% of WT and ~90% of TAX1BP1 KO cells. (Fig. 3C). Furthermore, upon puromycin washout, foci were cleared to a greater extent in TAX1BP1 high-expressing rescue lines in which only ~5-10% of cells retained aggregates compared to 30% in WT cells, and >70% in the parental TAX1BP1 KO (Fig. 3D), suggesting an active role for TAX1BP1 in promoting aggregate clearance.

Because we did not observe robust foci formation in the high-expression TAX1BP1-rescue lines (Fig. 3C, Supplementary Fig. 3B), we treated cells with Bafilomycin A during puromycin treatment and washout to block autophagy. In WT cells, Bafilomycin A resulted in increased aggregate retention after washout (Fig. 3E, 3F, larger field of view shown in Supplementary Fig 3C). However, in TAX1BP1 KO cells, Bafilomycin A barely exacerbated the defect in aggregate clearance, suggesting that the autophagy pathway is nonfunctional in the presence of proteotoxic stress (Fig. 3E, 3F). Bafilomycin A was sufficient to restore UB-positive foci in TAX1BP1 rescue cell lines (Fig. 3E, 3F, Supplementary Fig. 3C), resulting in UB-positive foci which colocalized with FLAG-TAX1BP1 and tended to loosely accumulate in the perinuclear region, similar to that observed upon addition of Bafilomycin A to WT cells (Fig. 3E, Supplementary Fig. 3D). Our data demonstrate that TAX1BP1-medated aggregate clearance is dependent upon expression level and capable of accelerating degradation of insoluble protein aggregates by promoting selective flux through the autophagy pathway.

TAX1BP1 contains several protein-interacting domains, including an N-terminal SKICH domain, at least 3 less studied coiled-coil regions, a canonical LIR motif and putative noncanonical CLIR motif, as well as two C-terminal zinc finger (ZF) UB-binding domains^24,25,27,28^. We created a variety of truncation and point mutants to test the requirements for these domains in TAX1BP1-mediated aggrephagy (Fig. 4A). Each mutant was stably expressed in the TAX1BP1 KO background and assessed for puromycin-induced aggregate formation and clearance. Aggregate formation occurred with minor variation similarly in WT and all mutant cell lines as well as a GFP-only control (Fig. 4Bi,ii, 4C, larger fields in Supplementary Fig. 4A). However, substantial differences were observed amongst the mutants in aggregate clearance. N-terminal SKICH deletion (ΔSKICH) as well as both the LIR (W49A) and CLIR (V143S) point mutants localized to UB-positive foci and rescued clearance to the same extent as WT TAX1BP1 (Fig. 4Biii, iv, v, 4D, Supplementary Fig. 4A). The structure of the TAX1BP1 SKICH domain shows that the canonical LIR (W49) is mostly buried within the hydrophobic interior of the folded SKICH domain and therefore is likely not a functional LIR domain^36^. A truncation mutant lacking both C-terminal ZF domains did not localize to puromycin-induced foci and was unable to rescue aggregate clearance (Fig. 4Bvi, 4D). To determine whether one or both ZF domains was essential, we created point mutants targeting both ZF domains (ZF1/ZF2: Q743A/E747K/ Q770A/E774K) as well as each domain individually (ZF1:Q743A/E747K, ZF2:Q770A/E774K). The double mutant, ZF1/ZF2, as well as the single ZF2 mutant, failed to localize to UB foci after puromycin treatment and both were unable to rescue aggregate clearance (Fig. 4Bviii, ix, 4D). In contrast, point mutation of ZF1 had little effect on TAX1BP1 localization and rescued aggregate clearance similarly to WT TAX1BP1 (Fig. 4Bvii, 4D). This is consistent with a prior report that only the ZF2 of TAX1BP1 is capable of binding UB^25^. An N- and C-terminally truncated mutant (ΔSKICH/ΔZF) did not localize to aggregates but instead formed aberrant clusters in the cytosol and was inferior to the ZF mutants in its ability to rescue clearance (Fig. 4Bx, 4D, Supplementary Fig. 4A).

**Figure 4.**
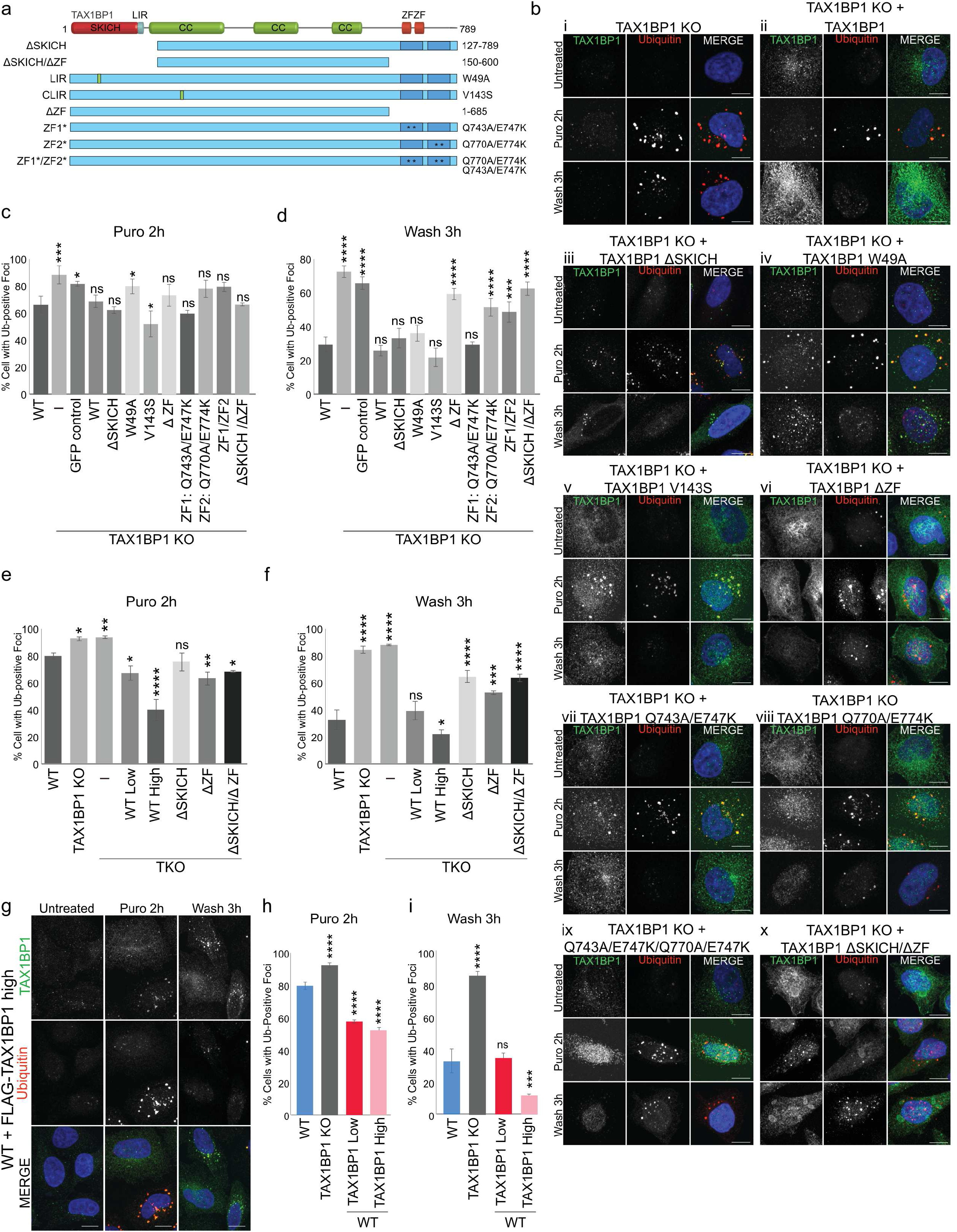
Requirements for TAX1BP1 domains in aggrephagy. **a,** TAX1BP1 truncation or point mutations used in this study. **b, i-x,** TAX1BP1 KO and TAX1BP1 KO with stable expression of TAX1BP1 mutants were exposed to 5 μg/ml puromycin for 2 h, after which cells were either fixed for imaging or washed and followed for a further 3 h in full media; scale bar 10μm, larger fields of view shown in Supplementary Figure 4A. **c,** Quantification of UB-foci formation observed in (b). **d,** Quantification of UB-foci clearance observed in (b), percent of cells containing UB-positive foci was assessed in ~ 200 cells per condition. **e,** Quantification of UB-foci formation observed in TKO (OPTN/NDP52/TAX1BP1). See Supplementary Figure 4C for images. **f,** Quantification of UB-foci clearance observed in TKO. See Supplementary Figure 4C for images, percent of cells containing UB-positive foci was assessed in ~ 200 cells per condition. **g,** WT cells stably expressing high levels of FLAG-TAX1BP1 were exposed to 5 μg/ml puromycin for 2 h, after which cells were either fixed for imaging or washed and followed for a further 3 h in full media; scale bar 10μm. **h,** Quantification of UB-foci formation observed in (g) and in Supplementary Figure 4D. **i,** Quantification of UB-foci clearance observed in (g) and in Supplementary Figure 4D, percent of cells containing UB-positive foci was assessed in ~ 200 cells per condition. All quantification is displayed as mean ± s.d. from 3 independent experiments using one-way ANOVA test comparing all to WT (\**P*<0.05, \*\**P*<0.01, \*\*\**P*<0.001, \*\*\*\**P*<0.0001) and Tukey’s post hoc test. The number of observations used in each experimental series and *P* values for all comparisons are included in Table 1. All images are representative of at least 3 independent experiments.

Very similar results were observed in triple knockout (TKO: TAX1BP1 KO, OPTN KO, NDP52 KO) cells (Supplementary Fig. 4B, 4C), ruling out potential redundancy that may mask requirements for TAX1BP1 domains through functionally related autophagy receptors. Low, near-endogenous expression of TAX1BP1 rescued aggregate clearance to that of WT cells while higher expression increased clearance compared to WT cells (Fig. 4E, 4F, Supplementary Fig. 4C). In contrast, neither the ΔSKICH, ΔZF nor ΔSKICH/ΔZF mutants were able to fully rescue aggregate clearance demonstrating the need for these protein-interaction and UB-binding domains (Fig. 4F, Supplementary Fig. 4C). Thus, full length TAX1BP1 was able to rescue aggregate clearance in the absence of NDP52 and OPTN, indicating that TAX1BP1 promotes aggrephagy (Fig. 4F).

We also examined whether TAX1BP1 overexpression in WT cells beyond the level of endogenous TAX1BP1 expression could promote aggregate clearance, thus addressing the therapeutic potential of TAX1BP1. TAX1BP1 protein expressed in WT cells behaved similarly to the endogenous protein, remaining diffuse in untreated cells and colocalizing with UB-stained foci upon puromycin treatment (Fig. 4G, Supplementary Fig. 4D). Aggregate formation upon 2h puromycin treatment was decreased in both low (~20% decrease) and high (~30% decrease) - overexpressing TAX1BP1 cells (Fig. 4H). Clearance of aggregates in WT cells with low overexpression of TAX1BP1 was similar to that in WT cells expressing only endogenous TAX1BP1 (Fig. 4I, Supplementary Fig. 4D), whereas, higher levels of TAX1BP1 expression increased clearance upon puromycin washout – only ~10% of cells retained UB-positive foci in contrast to more than 30% of WT cells (Fig. 4G, 4I).

Overexpression of WT TDP-43 and polyQ huntingtin fragments is cytotoxic in *in vitro* and *in vivo* model system^20,37–44^. Stimulation of autophagy increases clearance of WT and mutant TDP-43, huntingtin fragments and attenuates cytotoxicity^20,43,45–48^. Therefore, WT, TAX1BP1 KO, and high-expressing TAX1BP1-rescue cells were exposed to a variety of stresses, including low dose proteasome inhibition and expression of aggregation-prone proteins, such as the ALS-associated, EGFP-TDP-43, and varied length model substrates carrying expanded glutamine tracts (polyQ) expressed from exon 1 of the huntingtin-encoding gene (Supplementary Fig. 5A, 5B). Cells were either infected or transfected on day 1 and assessed daily for six days. Loss of TAX1BP1 decreased viability compared to WT cells upon exposure to each form of proteotoxic stress (Fig. 5A, Supplementary Fig. 5C). Expression of TAX1BP1 in the knockout line was able to partially restore cell viability (Fig. 5A, Supplementary Fig. 5C).

**Figure 5.**
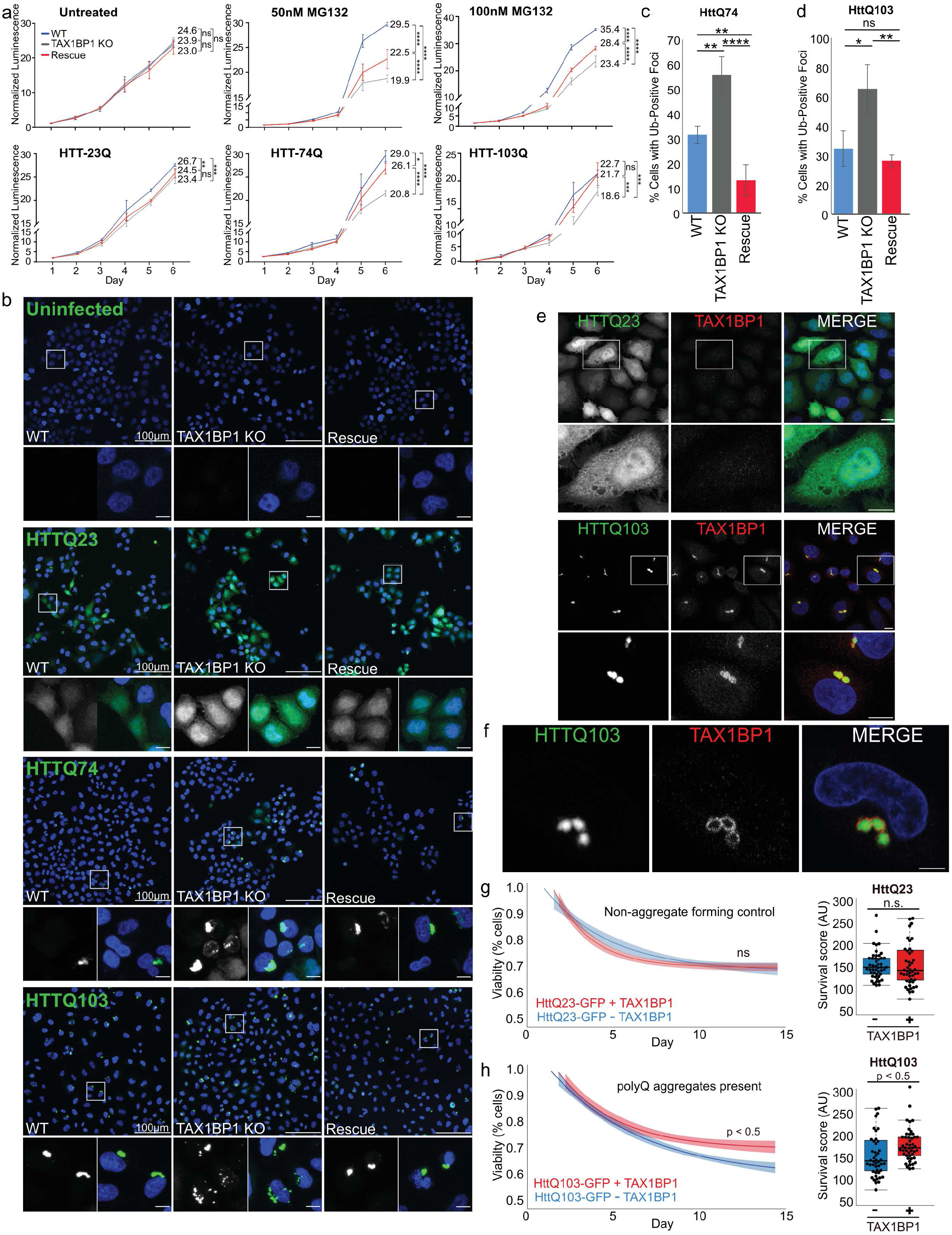
TAX1BP1 mediates aggrephagy of cytotoxic aggregation-prone proteins. **a,** WT, TAX1BP1 knockout, or TAX1BP1 knockout with stably-expressed TAX1BP1 rescue cell lines were exposed to proteotoxic stressors as indicated on Day 1, then followed for 6 days during which viability was measured by quantification of ATP production. Relative viability represents normalized luminescence displayed as mean ± s.d. from 3 independent experiments; significance was assessed using two-way ANOVA test (\*\*\*\**P*<0.0001, \*\*\**P*<0.001, \*\**P*<0.01, \**P*<0.05) with Tukey’s post hoc test. *P* values and normalized viability measurements shown on graphs are for day 6 comparisons. The individual measurements for each time point and conditions used in each experimental series and *P* values for all comparisons are included in Table 1. **b,** WT, TAX1BP1 knockout, or rescue cells uninfected or infected with virus expressing HttQ23-EGFP, HttQ74-EGFP, or HttQ103-EGFP were assessed 4 days post-infection. **c,** Quantification of HttQ74-EGFP aggregates observed in (b), **d,** Quantification of HttQ103-EGFP aggregates observed in (b), percent of cells containing GFP-positive foci was assessed in ~ 200 cells per condition in 3 independent experiments. Quantification is displayed as mean ± s.d. from 3 independent experiments using one-way ANOVA test (\*\**P*<0.01,\*\*\**P*<0.001,\*\*\*\**P*<0.0001) and Tukey’s post hoc test. The number of observations used in each experimental series is included in Table 1. All images are representative of at least 3 independent experiments. **e,** Immunofluorescence labeling of endogenous TAX1BP1 in cells infected with HttQ23-EGFP or HttQ103-EGFP. Maximum intensity projections shown. **f,** A single 1μm slice is shown from images taken of immunofluorescence labeling of endogenous TAX1BP1 in cells infected with HttQ103-EGFP as in **(e)**. Scale bars 10μm. **g, h,** Graphs show line fitted to the number of BFP-positive nuclei counted daily for iPSC-derived neurons with or without stable TAX1BP1 overexpression infected with HttQ23-EGFP (g) or HttQ103-EGFP (h). Ribbon represents 95% confidence interval around the fitted line. Beeswarm box plots compare survival scores determined by performing permutation analysis using the means of all slopes (center line = median, box limits = first to third quartile, whiskers = minimum and maximum). The number of observations used in each experimental series is included in Table 2.

To determine whether TAX1BP1 can aid in clearance of Htt-polyQ proteins, we infected WT, TAX1BP1 KO, and TAX1BP1-rescue cells with virus to express HttQ23-EGFP, HttQ74-EGFP and HttQ103-EGFP and assessed aggregate levels 4 days post-infection (Supplementary Fig. 5A). Due to the low number of glutamine repeats, HttQ23-EGFP does not form inclusions and therefore is a useful control when comparing to the aggregate-forming HttQ74-EGFP and HttQ103-EGFP proteins. HttQ23-EGFP expression was robust and as expected the protein remained largely diffuse and cytosolic in both WT and TAX1BP1 KO cells (Fig. 5B). In contrast, TAX1BP1-deficient cells exhibited increased focal aggregates of both Htt74Q-EGFP and Htt103Q-EGFP compared to WT cells (Fig. 5B, 5C, 5D). Rescue of the TAX1BP1 KO line with expression of FLAG-TAX1BP1 restored clearance of HttQ74-EGFP and Htt103Q-EGFP to that of WT cells, indicating that TAX1BP1 is highly effective in directing clearance of huntingtin polyQ proteins (Fig. 5B, 5C, 5D). Endogenous TAX1BP1 also colocalized with HttQ103-EGFP aggregates in WT cells (Fig. 5E) appearing to enclose aggregates in many instances (Fig. 5F, Supplementary Figure 5D).

Because TAX1BP1 overexpression can promote clearance of aggregates beyond that in WT cells and because we found TAX1BP1 highly and specifically-expressed in brain (Figure 2D, 2F), we examined whether TAX1BP1 could provide a protective effect in iPSC-derived neurons exposed to huntingtin proteins. iPSC-derived neurons with or without stable expression of TAX1BP1 were infected with virus expressing either the non-aggregating control construct, HttQ23-EGFP, (Figure 5G) or the aggregate-forming HttQ103-EGFP (Figure 5H) and imaged daily for 15 days to assess viability via nuclei count (NLS-BFP). The rates of cell death, assessed by comparing slopes obtained from lines fitted to cell number over time, did not differ between WT and TAX1BP1-overexpressing neurons infected with the non-aggregate-forming HttQ23-EGFP (Figure 5G). However, TAX1BP1 overexpression significantly improved survival in neurons exposed to the aggregate-forming HttQ103-GFP protein (Figure 5H), consistent with our observations in HeLa cells.

Here we report a broad role for TAX1BP1 in protein homeostasis. Our results demonstrate that loss of TAX1BP1 leads to decreased ability to target insoluble protein for degradation. Overexpression of TAX1BP1 further promotes aggregate clearance and rescues cell viability upon exposure to varied proteotoxic insults, including translation stress, proteasome inhibition, and exposure to aggregate-prone proteins such as TDP-43 and huntingtin-expanded polyQ model substrates (Supplementary Figure 5E).

Aggrephagy has potential to mitigate neurodegenerative proteinopathies. Focus on the proteins that provide specificity in targeting aggregates, such as TAX1BP1, may be valuable, as these are likely deciding factors in directing the autophagy response. Using single or combinatorial knockouts of OPTN, NDP52, TAX1BP1, NBR1, and p62, we observed distinct roles for these varied autophagy receptor proteins in aggrephagy. Though they may function *en masse* to maintain protein homeostasis, the individual proteins exhibit functional and spatial distinctions at the subcellular and tissue level. For example, autophagy receptor proteins are recruited independently to distinct microdomains surrounding bacteria and mitochondria where they perform nonredundant roles during xenophagy and mitophagy, respectively^26,49^. Furthermore, TAX1BP1 is robustly expressed in human brain lysate as well as primary rat cortical neurons, distinguishing it from NDP52. TAX1BP1 is also associated with the insoluble protein fraction in primary rat cortical neurons exposed to proteotoxic stress. Amongst the other autophagy receptor proteins, only p62 showed similar association with the insoluble fraction. TAX1BP1 and its paralog, NDP52, share similar domain structures; however, recent studies suggest structural differences in the organization of the SKICH domains as well as distinct ATG8 binding affinities^25,50^. TAX1BP1 is also more broadly conserved than NDP52 among mammals^25^, suggesting that TAX1BP1 fills nonredundant essential roles. Future studies examining TAX1BP1 expression levels during aging and in animal models of neurodegenerative disease are warranted.

TAX1BP1 recruitment to protein aggregates requires the C-terminal ubiquitin-binding domain and its function in promoting aggrephagy further necessitates the N-terminal SKICH domain. However, none of our TAX1BP1 mutants were completely dead in terms of rescue effect, suggesting that other cellular functions of TAX1BP1 may be involved. One such role is to act as a Myosin VI cargo adaptor protein for mediating autophagosome maturation, which could contribute to clearance dependent upon other upstream factors, thus explaining partial rescue^25^. Additionally, TAX1BP1 is best studied for its role in negatively regulating nuclear factor-kb (NF-κB) and interferon regulatory factor (IRF) 3 via an interaction with the deubiquitinase, A20, thus restricting pro-inflammatory signaling and immune response^51,52^. Though links between TAX1BP1’s role as an autophagy adaptor and the immune response are yet to be understood, it is well known that protein aggregation, inflammation, and autophagy are intertwined^53^. Studies have shown that exposure to disease-associated protein aggregates can elicit innate immune response in glial cells and that LPS-induced inflammation results in enhanced aggregate formation in disease models suggesting a synergistic relationship between proinflammatory response, proteostasis and neurodegeneration^54,55^. Autophagy may function to downregulate inflammatory signaling and TAX1BP1 may be an important link between detection and monitoring of cellular protein aggregates and the inflammatory response.

Notably, ectopic expression of TAX1BP1 in knockout cells was able to rescue aggregate clearance to WT levels and increased overexpression of TAX1BP1 in both the knockout and WT cells was able to reduce aggregate levels below those observed in WT cells. Furthermore, TAX1BP1 overexpression in IPSC-derived neurons was protective against huntingtin aggregate-induced toxicity. TAX1BP1 thus plays a general role in promoting aggrephagy and future studies aimed at increasing TAX1BP1 expression or stability *in vivo* present promising therapeutic potential in addressing proteinopathies. An increased understanding of targeting specificity of selective autophagy receptor proteins for protein aggregates may make autophagy-stimulating approaches more specific and effective in treatment of protein misfolding diseases.

## Acknowledgements

We thank Drs. Alicia Pickrell, Malavika Raman, Achim Werner, David Beck and Richa Lomash for critical reading of the manuscript. We also thank the NINDS Light Imaging Facility, the NICHD Imaging Facility, and the NHLBI Flow Cytometry Core Facility. We thank Dr. Harm Kampinga for sharing the EGFP-HttQ43 and EGFP-HTTQ74 plasmids, John Badger and Dr. Katherine Roche for providing the primary rat cortical neurons and expertise. This work was supported in part by the National Institutes of Health Grants NINDS intramural program (R.J.Y) and NIGMS Postdoctoral Research Associate Fellowship (S.A.S.).

## Author Contributions

S.A.S. and R.J.Y. conceived the project; S.A.S. and R.J.Y. designed experiments; S.A.S and H.V.S. performed experiments; G.K. performed image analysis for neuron viability; M.E.W. provided iPSCs and relevant expertise, S.A.S and R.J.Y. wrote the manuscript and all authors contributed to editing the manuscript.

## Methods

### Cell Culture and reagents

HeLa and HEK293T were cultured in DMEM (Life Technologies) supplemented with 10% (v/v) FBS (Gemini Bio Products), 10 mM HEPES (Life Technologies), 1 mM sodium pyruvate (Life Technologies), 1mM non-essential amino acids (Life Technologies) and 2 mM glutamine (Life Technologies). HeLa cells were acquired from the ATCC and authenticated at the Johns Hopkins GRCF Fragment Analysis Facility using STR profiling. Testing for mycoplasma contamination was performed bimonthly using the PlasmoTest kit (InvivoGen). Plasmids were transfected using either X-tremeGENE 9 (Roche), polyethylenimine (Polysciences) or Avalanche-OMNI (EZ Bio-systems). Primary cultured cortical neurons were isolated from male and female embryonic day 18 Sprague-Dawley rats and maintained in Neurobasal medium (Gibco) supplemented with 2mM glutamine (Life Technologies) and B-27 Supplement (Gibco). All rat procedures were performed according to a protocol (#1171) approved by the National Institutes of Health NINDS Institutional Animal Care and Use Committee. Primary neurons were treated with MG132 on day-in-vitro (div) 10.

For puromycin (Invivogen), MG132 (Sigma-Aldrich) or Bafilomycin A (Sigma-Aldrich) treatments, cells were treated with the indicated concentrations of drug in full media and either harvested as described or washed three times in full media and returned to full media for the indicated durations.

### Antibodies

Rabbit mono- and polyclonal antibodies used for immunoblotting include: NDP52 (CST, 60732S), TAX1BP1 (CST, 5105S), Actin (CST, 4967S); GAPDH (Sigma, G9545-200UL), OPTN (Proteintech, 10837-I-AP) and TAX1BP1 (Sigma, HPA024432); rabbit antibodies used for immunofluorescence include TAX1BP1 (Sigma, HPA024432). Mouse Monoclonal antibodies used for immunoblotting were: GFP (Roche, 11814460001), ubiquitin (Millipore, MAB1510), NBR1 (Abnova, H00004077-M01) and p62 (Abnova, H00008878-M01); mouse antibodies used for immunofluorescence include ubiquitin FK2 (Biomol International, PW8810-0500). Secondary AlexaFluor^®^ (ThermoFisher) conjugated antibodies were incubated for 1h RT at 1:1000. DAPI counterstain (ThermoFisher) at 300nM was incubated for 10 min RT.

### TALEN and CRISPR gene knockout cell lines

Construction of OPTN, NDP52, TAX1BP1, TKO and pentaKO knockout HeLa lines were reported previously ^17^. All knockout lines were generated using CRISPR guide RNAs (gRNAs) chosen to target one or more exons common to all splicing variants of the gene of interest (all information listed in Supplementary Table S1). Oligonucleotides (Operon) containing CRISPR target sequences were annealed and ligated into the linearized gRNA_Cloning vector which was a gift from George Church (Addgene #41824) or SpCas9-2A-Puro, which was a gift from Feng Zhang (Addgene #48139). HeLa cells were cotransfected using XtremeGENE9 (Roche) with either the the gRNA_Cloning vector, pCDNA YFP-C1, and the hCas9 plasmid, which was a gift from George Church (Addgene #41815) or with SpCas9-2A-Puro. Forty-eight hours post-transfection, YFP-positive cells were isolated either via fluorescence activated cell sorting or puromycin selection and serially diluted for single colony clones. Single colonies were expanded and screened for depletion of the targeted gene product by immunoblotting. DNA was extracted using the Zymo gDNA Isolation Kit and genotyped using primers targeting fragments of the genomic DNA from knockout clones containing the putative cleavage site (primers and sequences in Supplementary Table 1).

### Cloning and generation of stably infected cell lines

The Gateway Cloning (Life Technologies) system was used to generate pHAGE-GFP-TAX1BP1, pHAGE-N-FLAG-HA-TAX1BP1, and pHAGE-GFP-TDP43. Briefly, genes of interest were cloned into pDONR223 then recombined into destination vectors pHAGE-N-FLAG-HA or pHAGE-N-GFP using L Recombinase (Life Technologies) as per the manufacturer’s protocol. Mutations in cDNA sequences were introduced using PCR site-directed mutagenesis in the pDONR223 vector. All constructs generated in this study were verified by sequencing. Original plasmids containing EGFP-HttQ43 and EGFP-HttQ74 were kindly provided by Dr. Harm Kampinga (Groningen, Netherlands). pYES2/103Q was a gift from Michael Sherman (Addgene plasmid #1385) and pEGFP-Q23 was a gift from David Rubinzstein (Addgene plasmid #40261). All huntingtin protein constructs were cloned into pLEX-EF1α-EGFP for lentiviral expression. pDONR-TDP-43 WT YFP was a gift from Aaron Gitler (Addgene plasmid #27470). Primer sequences are available upon request.

For generation of stable cell lines, lentiviruses (pHAGE-vectors) were packaged in HEK293T cells. HeLa cells were transduced with virus for 24 hours with 8 μg/ml polybrene then selected for protein expression by drug-resistance (puromycin or blasticidin) or fluorescence activated cell sorting. Generation of low- and high-expressing cells was done via infection with various dilutions of virus and expression assessed via comparison to endogenous protein levels on immunoblot.

### Aggregate formation and clearance studies

For acute treatments, HeLa cells were grown on poly-D-lysine-coated (Sigma-Aldrich, P7280) coverslips and treated with either 5 μg/ml of puromycin (Invivogen) for 2 h or 1 μM MG132 (Sigma-Aldrich) for 8 h at 37°C. Cells were either fixed for imaging or assessed for clearance. For clearance, cells were washed three times in DMEM–10% FBS and released into drug-free medium for 3 h at 37°C. For long treatments, HeLa cells were grown on poly-D-lysine-coated coverslips and treated with 1 μg/ml of puromycin or 1 μM MG132 for 18 h at 37°C. Coverslips were fixed and processed for microscopy, as outlined above.

For huntingtin poly-Q-protein clearance assays, HeLa cells were seeded on poly-D-lysine-coated coverslips and transduced the following day with lentivirus expressing either HttQ23-EGFP, HttQ74-EGFP or HttQ103-EGFP for 24 hours with 8 μg/ml polybrene. Cells were fixed for imaging and assessed 4 days post-infection.

### Subcellular fractionation

Cells were washed once and scraped in cold PBS, pelleted, and lysed in RIPA buffer (Thermofisher, 89900) supplemented with Complete Protease Inhibitor Cocktail (Roche), 1mM EGTA, 1mM EDTA, 100mM chloroacetamide (Sigma-Aldrich) and 100mM DTT for 15 minutes at 4°C with end over end rotation. Five to ten percent of the total volume was collected, mixed with 4X LDS (Life Technologies), boiled, and reserved for the input fraction. RIPA-soluble fraction was obtained after centrifugation at 20,000 rpm for 15 min at 4°C, 4X LDS was added, and samples were heated to 99°C with shaking for 10 minutes. The insoluble fraction consisting of the remaining pellet was washed once with lysis buffer, spun at 20,000 rpm for 10 min at 4°C, then lysed in 1X LDS, 100mM DTT in PBS and heated to 99°C with shaking for 15 minutes. Protein concentration was determined via DC Protein Assay (BioRad). Soluble and insoluble fractions were normalized to the protein concentration of the input fraction. For analysis, 10-15% of RIPA-soluble fraction and 2.5-5% of insoluble fraction were loaded on 4-12% Bis-Tris gels and run using MOPS buffer (Life Technologies) and visualized by western blotting on PVDF. Western blotting was performed by wet transfer method in either NuPage transfer buffer or Tris-glycine transfer buffer (Life Technologies). Proteins were detected using horseradish peroxidase-coupled secondary antibodies (GE Healthcare Life Sciences, Piscataway, NJ) and ECL Plus or ECL Prime western blotting detection reagents (GE Healthcare Life Sciences). Images were acquired using an MP gel documentation system (Bio-Rad Laboratories). Quantification of immunoprecipitation bands was performed using the volume tools in Image Lab software (Bio-Rad Laboratories).

### Immunoblotting

After the indicated treatments, cells were harvested by scraping in cold PBS and either fractionated as described above or lysed in 2% SDS/1X PBS, heated to 99°C with shaking for 10 minutes, and spun at 20,000 rpm for 15 min at RT. Protein concentration was determined using DC Protein Assay (BioRad) and 20-50 ug of protein per sample was separated on 4–12% Bis-Tris gels using MOPS or MES running buffer (Life Technologies). Western blotting was performed by wet transfer method in either NuPage transfer buffer or Tris-glycine transfer buffer (Life Technologies). Proteins were detected using horseradish peroxidase-coupled secondary antibodies (GE Healthcare Life Sciences, Piscataway, NJ) and ECL Plus or ECL Prime western blotting detection reagents (GE Healthcare Life Sciences). Images were acquired using an MP gel documentation system (Bio-Rad Laboratories). Quantification of immunoprecipitation bands was performed using the volume tools in Image Lab software (Bio-Rad Laboratories). Human tissue panel blot was purchased (NOVUS Biologicals).

### Immunofluorescence Microscopy

HeLa cells were seeded on poly-D-lysine-coated (Sigma-Aldrich, P7280) coverslips and treated as indicated. Following treatment, cells were fixed at room temperature in 4% paraformaldehyde for 15 minutes, permeabilized and blocked with filtered IF buffer (0.1% Triton X-100, 3% goat serum, 1X PBS) for 1 hour at RT. For immunostaining, cells were incubated with indicated antibodies diluted in IF buffer overnight at 4°C, washed 3 times with IF buffer and incubated with Alexa Fluor-conjugated secondary antibodies (Life Technologies) 1:1000 in IF buffer for 1 hour at RT. Cells were then incubated with DAPI (Sigma) diluted in IF buffer at 1:10,000 for 10 minutes at RT. Cells were washed 2 times with IF buffer and 1 time with 1X PBS. Coverslips were mounted on slides using Prolong Gold Antifade (Life Technologies). Representative images were collected with an inverted laser scanning LSM 880 microscope (Carl Zeiss) using a 63X/1.4 objective Plan-Apochromat (Carl Zeiss). Images were collected as z-stacks captured at optimal thickness. Representative images shown are maximum intensity projections unless otherwise noted.

### Cell viability and Growth curve

For HeLa cell viability assays, cells were seeded in quadruplicate in white clear-bottom 96-well plates (Corning). The next day, cells were treated as indicated in figure legends. Cell viability was measured by using the Cell Titer-Glo system (Promega) according to the manufacturer's instructions. Values were normalized to the value for untreated samples for each cell line. Means and standard deviations were calculated, and statistical significance was calculated using one-way analysis of variance (ANOVA). One plate was measured each day using a Synergy H1Hybrid Multi-Mode Reader (Biottek).

### Statistical analysis for aggregate formation and clearance assays

For quantification, cells with puromycin-, MG132-, or HTT-induced foci were either counted manually in a blinded fashion or using the particle analysis plug-in in ImageJ. Macros created are available on request. At least 200 cells per sample were analyzed in three biological replicate studies. The number of observations used in each experimental series is included in Table 1. Means and standard deviations were calculated, and statistical significance was assessed. For comparisons between three or more groups, a one or two way-ANOVA with Tukey;s post-hoc analysis was performed using Prism V7 (GraphPad Software) as noted in the figure legend. Error bars represent standard deviation (SD): \**P*< 05, \*\**P*<.01, \*\*\**P*<.001, \*\*\*\**P*<.0001. The number of observations used in each experimental series was included in the figure legend or can be found in Table 1. All test statistics (e.g. F), degrees of freedom and *P* values are included in Table 1. Three biological replicates were performed for each experiment. To determine number of foci per individual cell, CellProfiler^56^, version 2.2.0 was used. All CellProfiler pipelines and detailed settings to reproduce the image analysis procedures are available upon request.

### Neuron Culture and Viability Assay

Induced pluripotent stem cells (iPSCs), parental line WTC11, were maintained in Essential 8 Flex medium (Gibco, A28583-01) supplemented with Essential 8 Flex Supplement (Gibco, A28584-01). iPSCs were grown on Matrigel-coated plates (Corning, 354277) and cell dissociation was performed using Accutase (StemCell Technologies, 07920). Stable expression of NLS-BFP (used for nuclei counting) was obtained by infecting cells with lentivirus expressing U6-NLS-BFP followed by sorting for BFP-positive cells. Stable expression of TAX1BP1 was obtained by infecting NLS-BFP-expressing cells with lentivirus expressing EF1α-FLAG-TAX1BP1 followed by puromycin selection.

For the viability assay, iPSCs grown in 6-well plates were infected with lentivirus expressing EF1α-HttQ23-EGFP or EF1α-HttQ103-EGFP, the following day, media was changed, and cells were allowed to recover for 1 day. Two days post-infection, cells were plated in induction medium: DMEM/F12 with HEPES (Gibco, 11330032) containing N2 Supplement (100X) (Gibco, 17502048), non-essential amino acids (100x) (Gibco, 11140050), glutamax (100X) (Gibco, 35050061), ROCK inhibitor Y-27632 (10mM) (Tocris, 1254), and doxycycline (2mg/ml) (Sigma, D9891). Induction medium was changed daily for 2 days. On day 3 post-induction, cells were dissociated using Accutase, counted, and plated at 50,000 cells/well in poly-L-ornithine-coated (PLO) (Sigma, P3655) 96-well plates (Perkin Elmer, 6055302) in BrainPhys neuronal medium (StemCell Technologies, 05790) containing B27 supplement (50X) (Gibco, 17504044), BDNF (10μg/ml) (PeproTech, 450-02), NT-3 (10μg/ml) (PeproTech, 450-03), and laminin (1μg/ml) (Gibco, 23017015). Half of the well volume was removed and replaced with fresh supplemented BrainPhys media every 3 days for the duration of the assay. iPSCs-derived neurons were imaged daily for 20 days. Sixteen fields of view were imaged per each well of three independent biological replicates. Cells were imaged using a 20X air objective (NA 0.75) with 1.5X optical zoom on a Nikon Ti-2 CSU-W1 spinning disk system with a photometrics 95B camera operated by Nikon Elements software equipped with temperature regulation and CO_2_ control.

### Statistical Analysis for Neuron Viability Assay

To determine iPCS-derived neuron survival, NLS-BFP-expressing nuclei were counted automatically using the R EBImage package^57^. Three biological replicates per condition were pooled together and the data were fitted to an exponential decay model using the R DRC package^58^ and the formula: (f(x) = c + (d-c)(exp(-x/e))) where (e) is the slope and (d) and (c) represent the upper and the lower limits around the slope. Outliers were detected and removed using interquartile range using the R Outliers package^59^. The survival score was calculated from the fitted model, in which the slope (e) represents the survival score of each population of cells counted at every time point for the designated treatment. For statistical comparison, a permutation test (a.k.a randomization test) was used. In brief, the delta mean score of the groups was compared to random delta mean scores of shuffled groups iterated 10,000 times. *P* value was determined by calculating the number of times the delta score was higher in the shuffled group than in the ground true group^58^. The number of observations used in each experimental series is included in Table 2. Code is provided in Supplementary Information.

### Data Availability

The datasets generated for all microscopy cell counting experiments are available as supplementary files and noted in the associated figure legends. The datasets generated and analyzed to assess neuron viability are available from the corresponding author on request. Associated code is available as supplementary files. Figures 1D, 1F, 1J, 3B, 3D, 3F, 4C, 4D, 4E, 4F, 4H, 4I, 5A, 5C, 5D, 5G, 5H, and Supplementary Figure 5C have associated raw data included in Tables 1 and 2. There are no restrictions on data availability.

### Code Availability

Code used to generate datasets and analyze neuron viability is available as supplementary files. All macros created for use with ImageJ/Fiji are available on request. CellProfiler pipelines and detailed settings to reproduce the image analysis procedures are available upon request.

## Supplementary Figure Legends

**Supplementary Figure 1.**
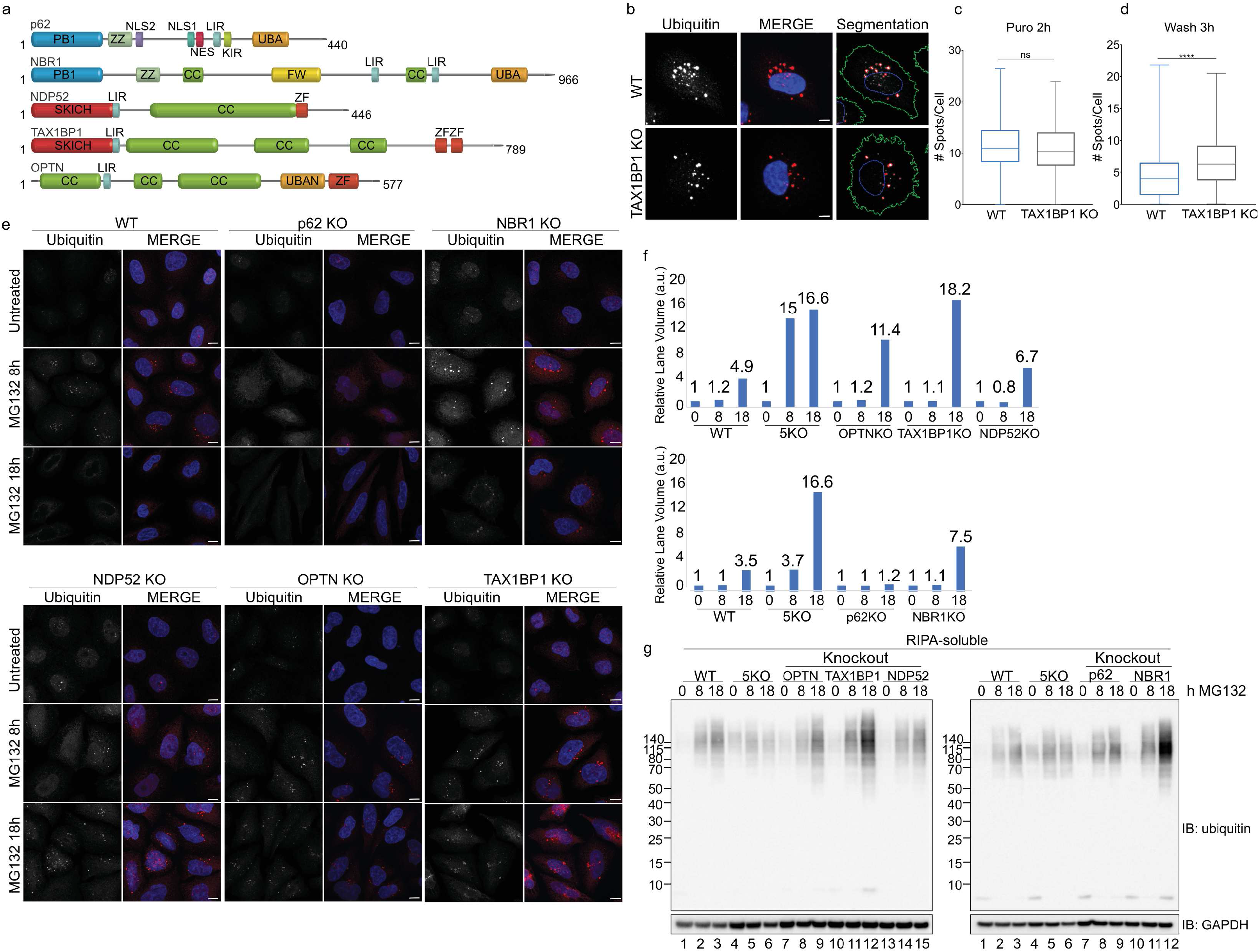
**a,** Schematic illustration of protein domain architectures of mammalian autophagy receptors OPTN, NDP52, TAX1BP1, p62, and NBR1. PB1, Phox and Bem1 domain; ZZ, ZZ-type zinc finger domain; NLS1 and NLS2, nuclear localization signals 1 and 2; NES, nuclear export signal; LIR, LC3-interacting region; KIR, Keap-interacting region; UBA, ubiquitin-associated domain; CC, coiled-coil domain; FW, four tryptophan domain; SKICH, SKIP carboxyl homology domain; ZF, Zinc-finger domain; UBAN ubiquitin binding in ABIN and NEMO domain. The size of the receptors (in number of amino acids) is indicated. **b,** Representative image of segmentation analysis performed using CellProfiler. **c, d,** Quantification of (b) using CellProfiler: number of foci per cell in WT or TAX1BP1 KO cells in 3 independent experiments at (c) 2 h puromycin or (d) 2 h puromycin followed by 3 h washout (For box plots, center line = median, box limits = first to third quartile, whiskers = minimum and maximum). **e,** WT and individual knockouts for p62, NBR1, NDP52, OPTN, and TAX1BP1 cell lines were exposed to 1 μM MG132 for 8 or 18 h, after which cells were either fixed for imaging or washed and followed for a further 3 h in full media; scale bar 10 μm. **f,** Quantification of Figure 1H determined by densitometry and normalized first to soluble GAPDH and subsequently to WT levels within each fraction. **g,** WT and individual KO lines for each autophagy receptor were treated with 1 μM MG132 for 8 or 18 h, fractionated into RIPA-soluble or -insoluble fractions and immunoblotted for total UB. Soluble fractions shown here, insoluble fractions shown in Figure 1H. All blots and microscopy images are representative of at least 3 independent experiments.

**Supplementary Figure 2.**
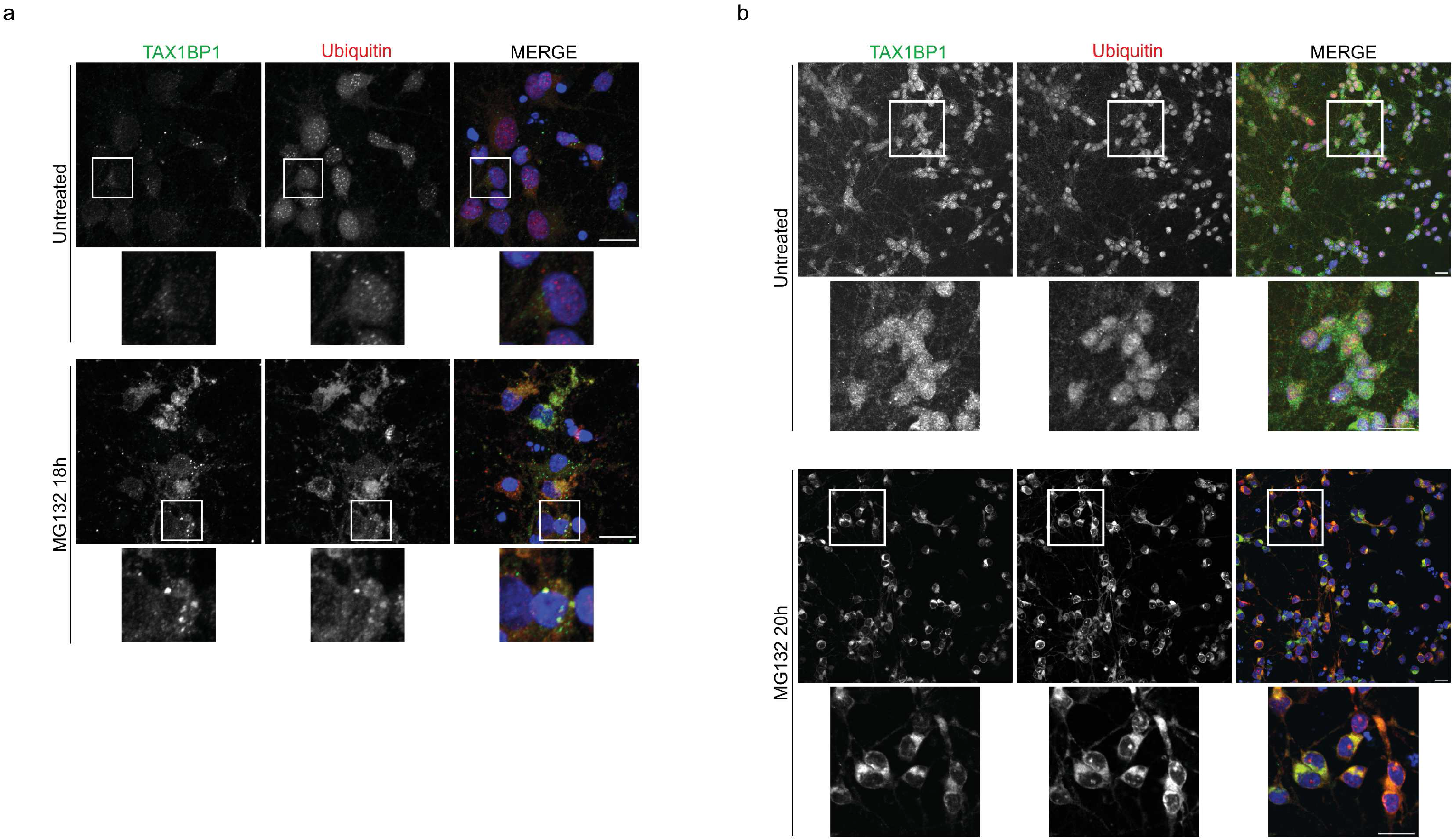
**a,** Primary rat cortical neurons treated with 1 μM MG132 for 18 h, after which cells were fixed for imaging and stained with antibodies for TAX1BP1 and UB; scale bar 10μm. **b,** Neurons derived from human induced pluripotent stem cells (iPSCs) treated with 1 μM MG132 for 20 h, fixed and stained with antibodies targeting TAX1BP1 and UB; scale bar 20 μm. All images are representative of at least 3 independent experiments.

**Supplementary Figure 3.**
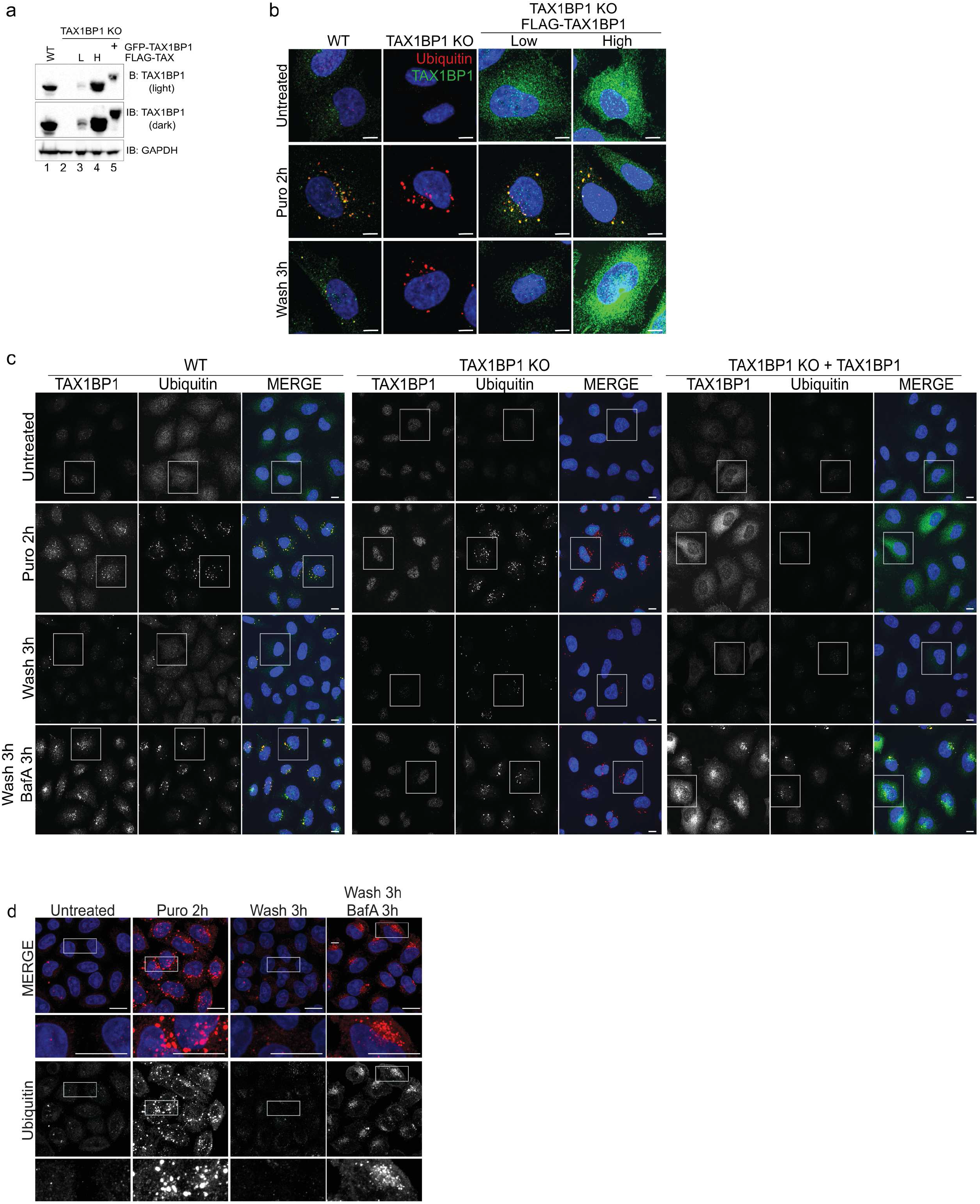
**a,** GFP- or FLAG-tagged TAX1BP1 was stably reintroduced into TAX1BP1 KO cells via viral infection. TAX1BP1 expression levels were titered for use in rescue experiments: L = low expression, H = high expression. **b,** Cell lines in (a) were exposed to 5 μg/ml puromycin for 2 h, after which cells were either fixed for imaging or washed and followed for a further 3 h in full media; scale bar 10μm. **c,** Full field of view images associated with Figure 3E, F showing WT, TAX1BP1 KO, or TAX1BP1 KO + FLAG-TAX1BP1 (H) cell lines exposed to 5 μg/ml puromycin in the presence or absence of 100 nM Bafilomycin A, after which cells were either fixed for imaging or washed and followed for a further 3 h in full media or in media containing Bafilomycin A. **d,** WT cells exposed to 5 μg/ml puromycin for 2 h in the presence or absence of Bafilomycin A, after which cells were either fixed for imaging or washed and followed for a further 3 h in full media with or without Bafilomycin A; scale bar 10μm. All images are representative of at least 3 independent experiments.

**Supplementary Figure 4.**
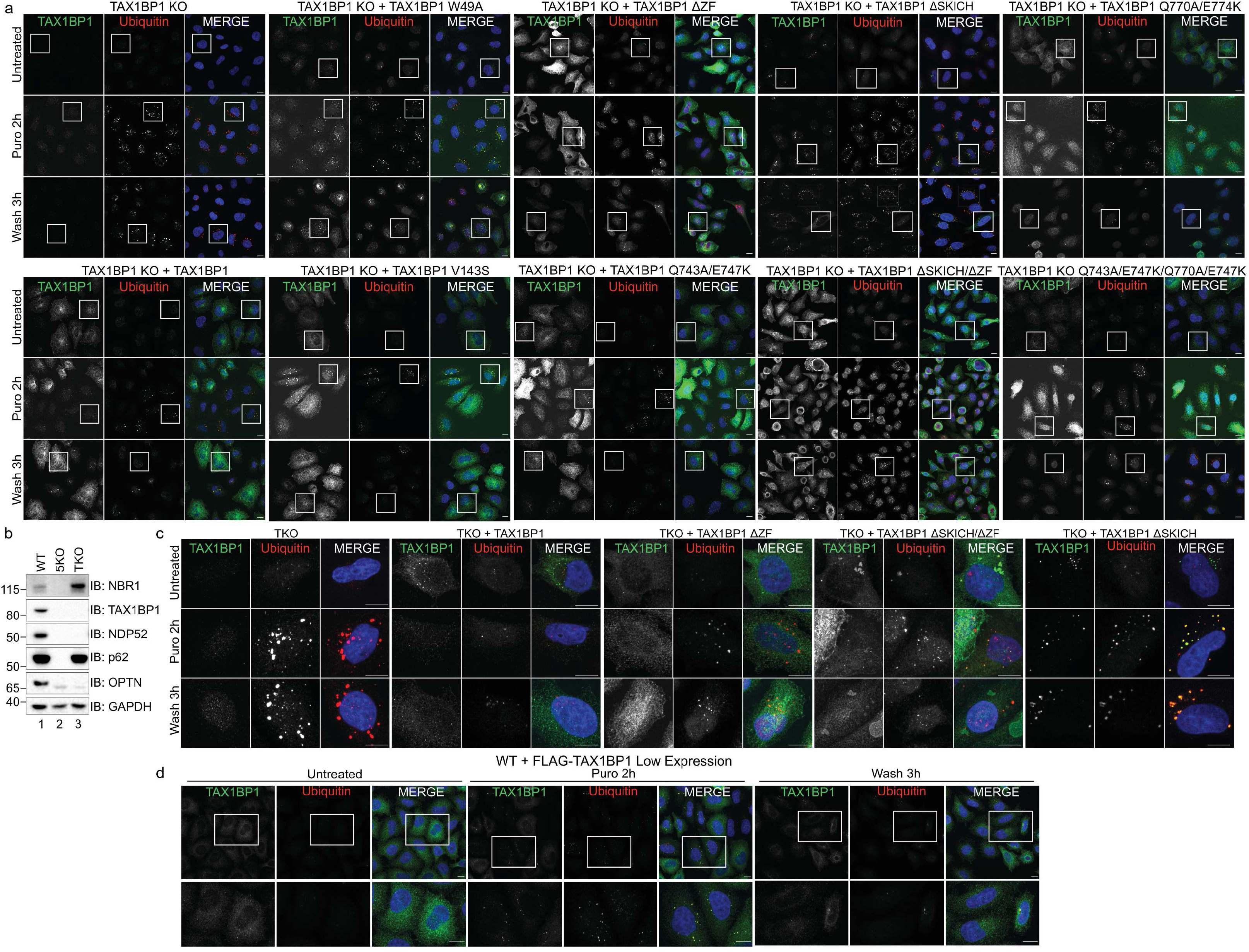
**a,** Full field of view images of all TAX1BP1 stable mutant expression cell lines exposed to 5 μg/ml puromycin for 2 h, after which cells were either fixed for imaging or washed and followed for a further 3 h in full media. Associated with Figure 4B, C, D. **b,** Validation of knockout cell lines. **c,** TKO (triple knockout: TAX1BP1, OPTN, NDP52) cell line with stable expression of TAX1BP1 mutants exposed to 5 μg/ml puromycin after which cells were either fixed for imaging or washed and followed for a further 3 h in full media; scale bar 10μm. **d,** WT cells stably expressing low levels of FLAG-TAX1BP1 were exposed to 5 μg/ml puromycin for 2 h, after which cells were either fixed for imaging or washed and followed for a further 3 h in full media; scale bar 10μm. All images are representative of at least 3 independent experiments.

**Supplementary Figure 5.**
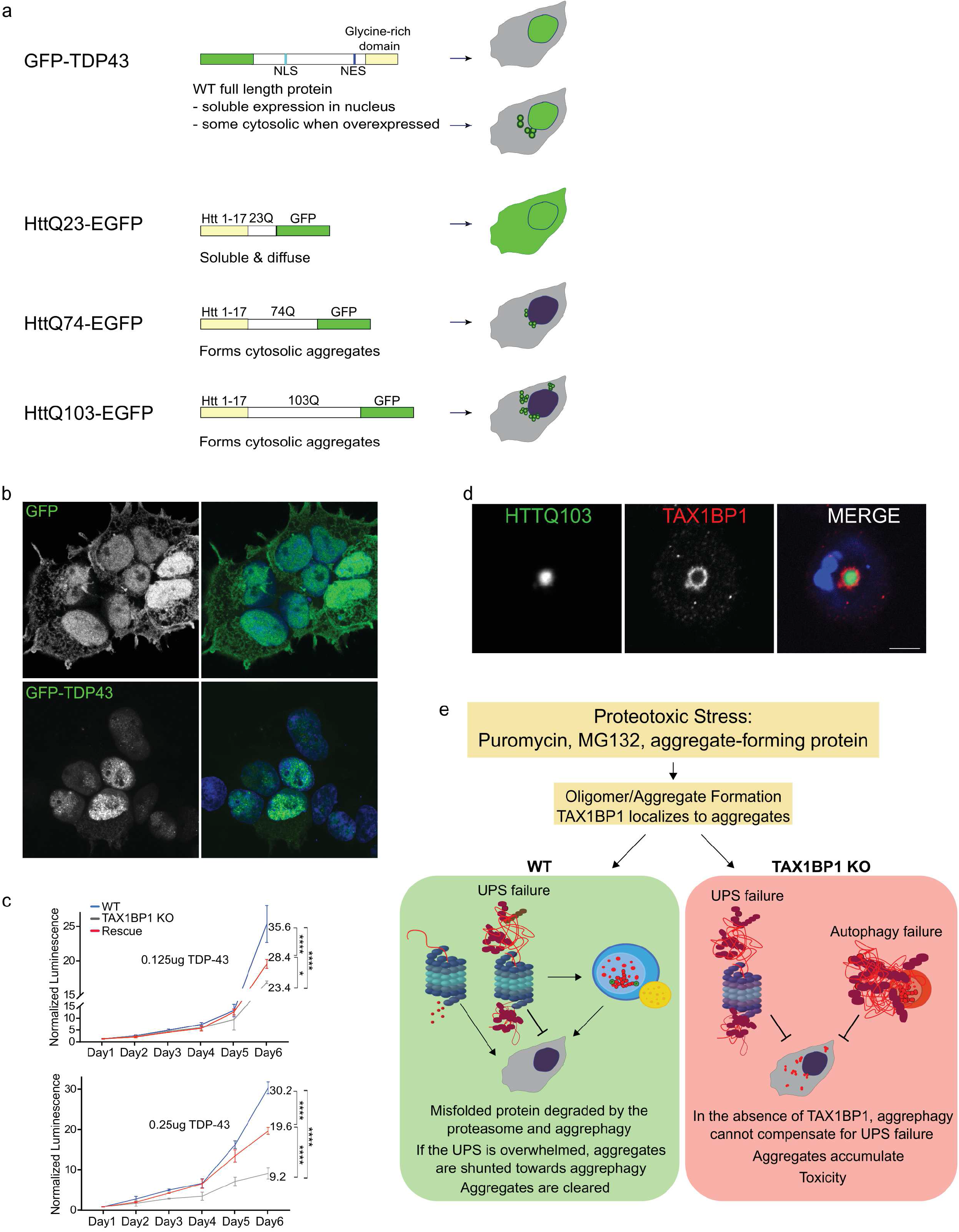
**a,** Constructs used in this study. **b,** Expression of GFP control or GFP-TDP-43. **c,** WT, TAX1BP1 knockout, or TAX1BP1-rescue cells transfected with DNA expressing EGFP-TDP-43 at the indicated concentrations on Day 1, then followed for 6 days during which viability was measured by quantification of ATP production. Relative viability represents normalized luminescence displayed as mean ± s.d. from 3 independent experiments; significance was assessed using two-way ANOVA test (\*\*\*\**P*<0.0001, \*\*\**P*<0.001, \*\**P*<0.01) with Tukey’s post hoc test. *P* values and normalized viability measurements shown on graphs are for day 6 comparisons. The individual measurements for each time point and conditions used in each experimental series and *P* values for all comparisons are included in Table 1. **d,** A single 1μm slice is shown from images taken of immunofluorescence labeling of endogenous TAX1BP1 in cells infected with HttQ103-EGFP; scale bar 10μm. **e,** Proteotoxic stress, induced by translational stress, proteasome inhibition, or expression of aggregate-promoting proteins causes misfolded or damaged proteins to assemble into toxic oligomers or aggregates. In WT cells (green panel), the proteasome and aggrephagy both work to remove potentially toxic protein products. If the proteasome is overwhelmed, aggregated protein is shunted to the autophagy pathway. In the absence of TAX1BP1 (red panel), aggrephagy is deficient - once the proteasome has become overwhelmed by misfolded or aggregated protein, there is decreased backup clearance via aggrephagy, and insoluble protein accumulates, leading to toxicity and cell death.

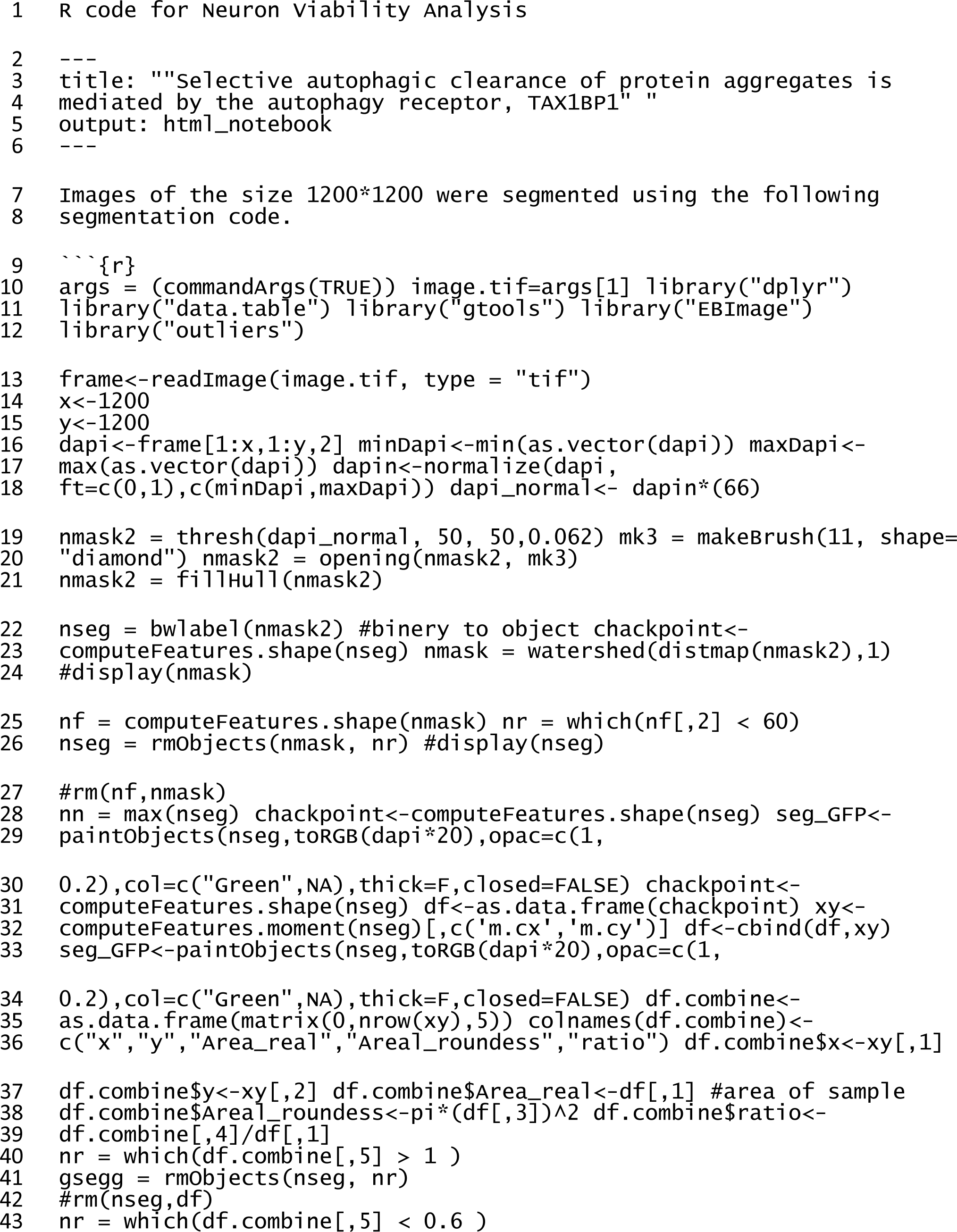
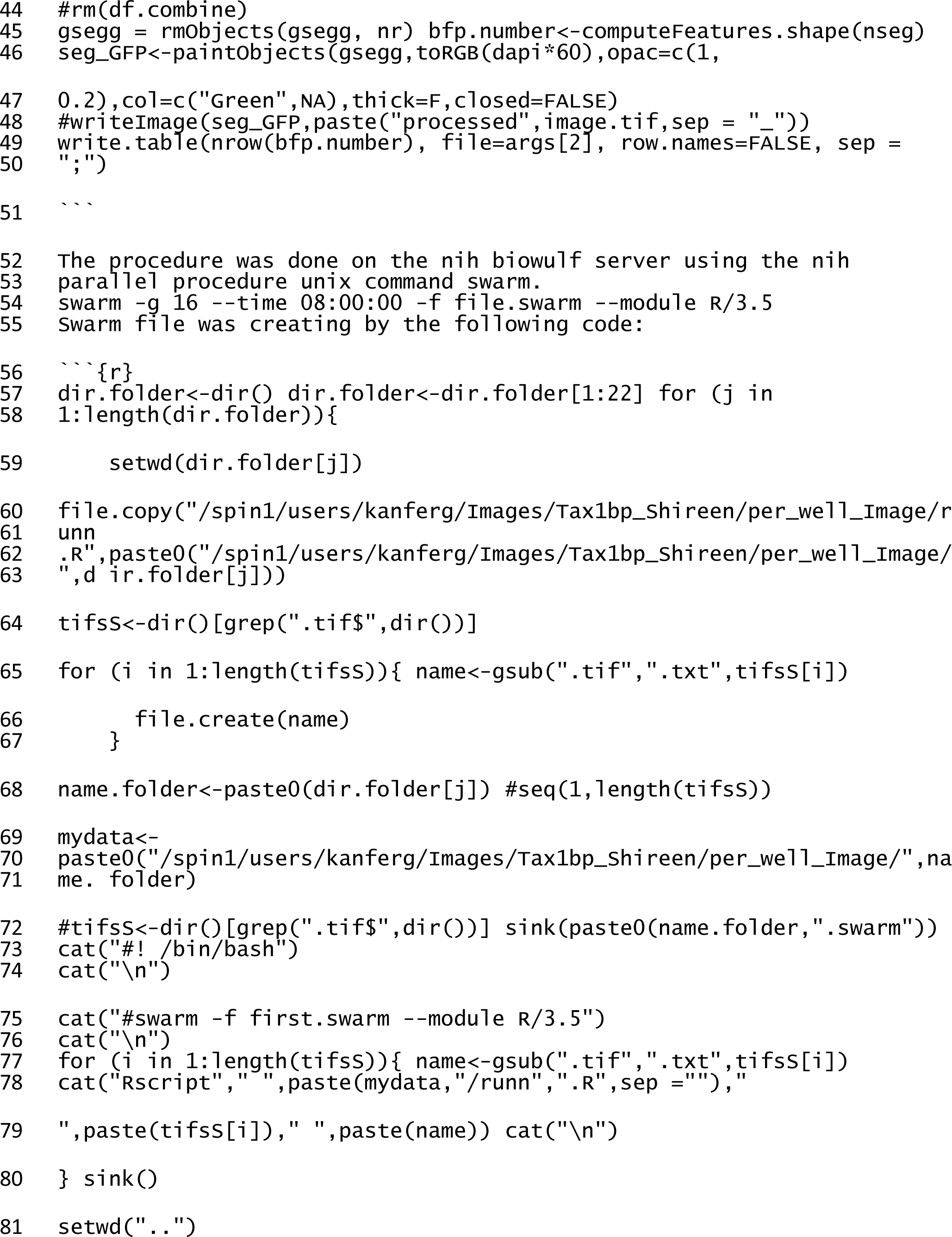
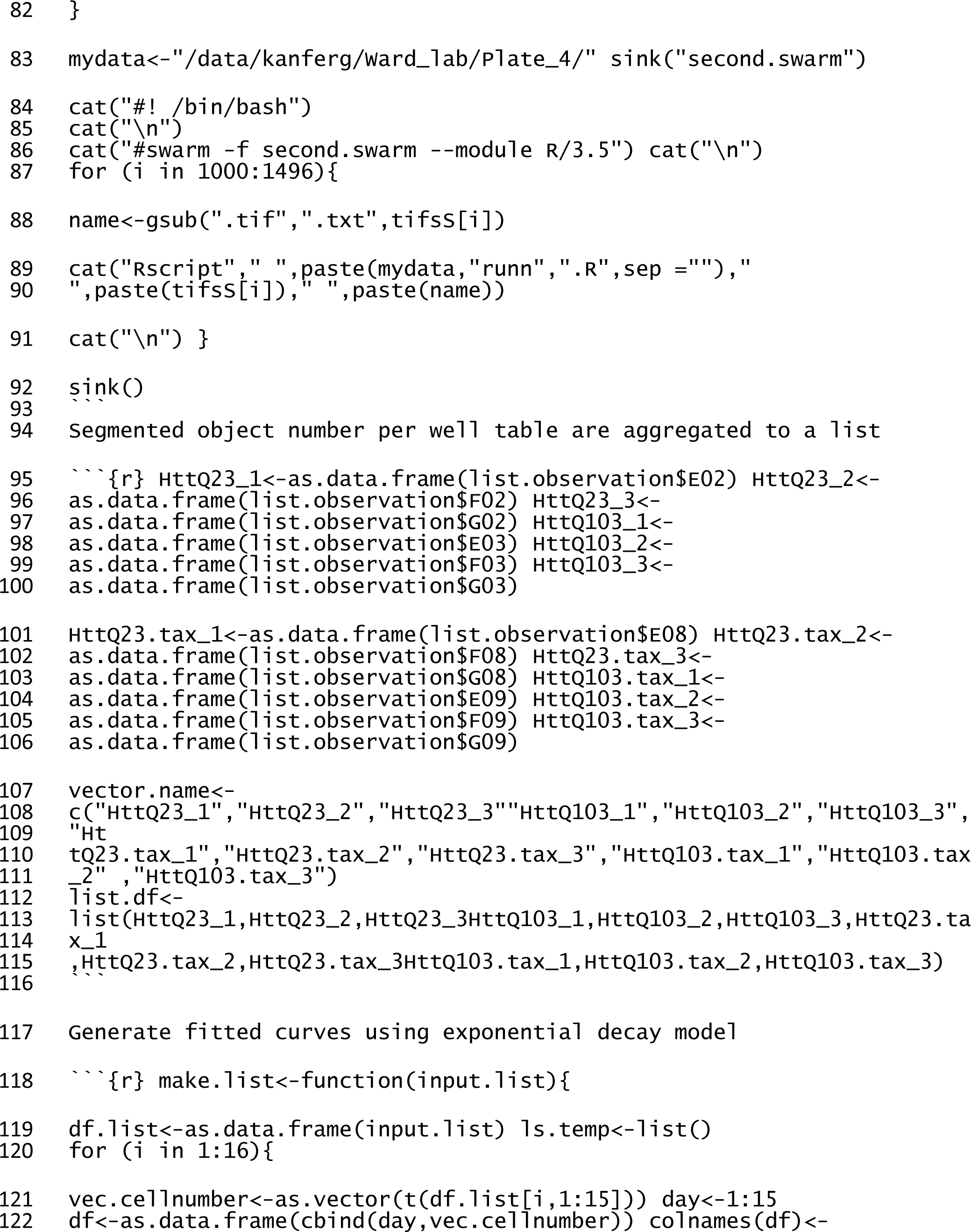
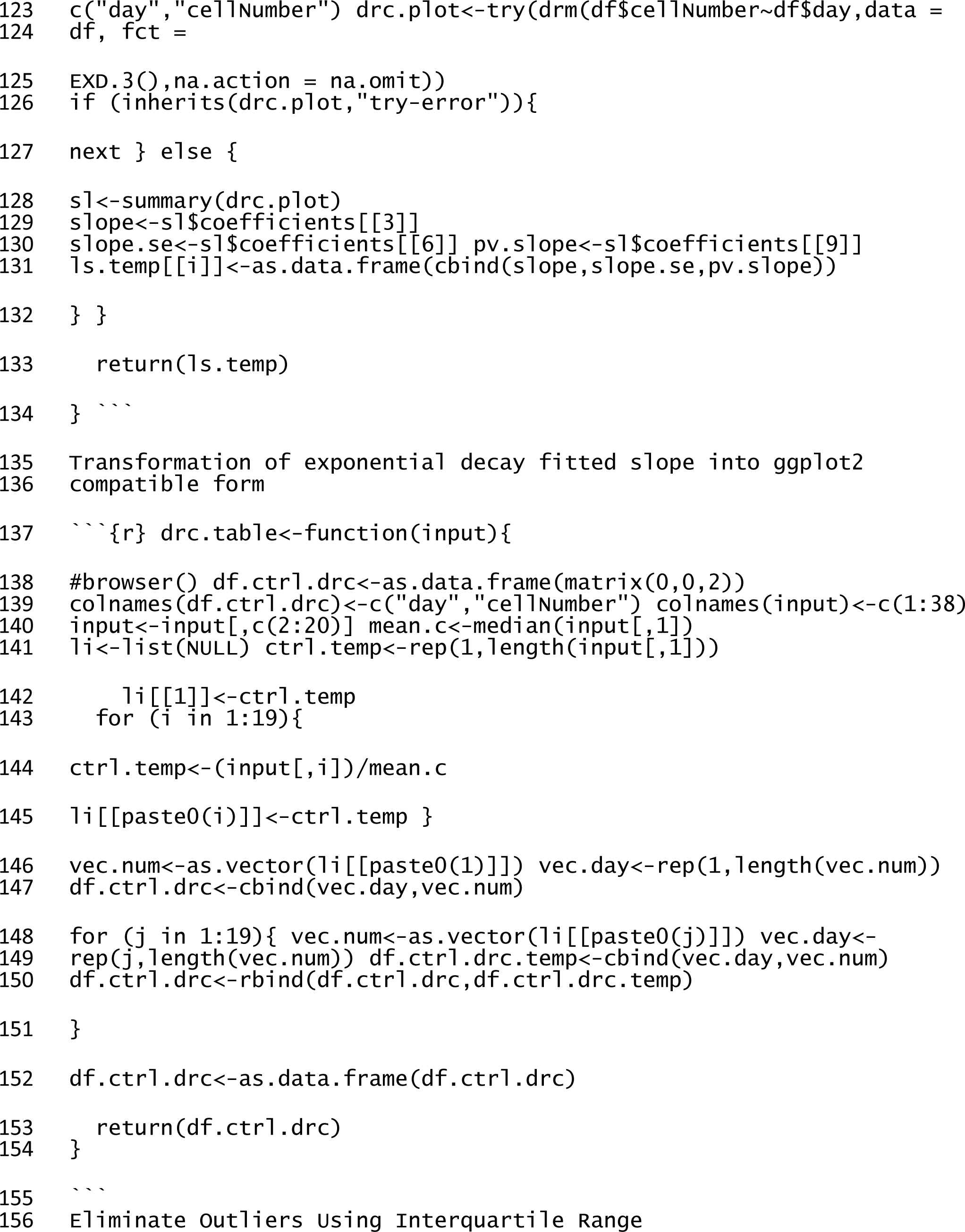
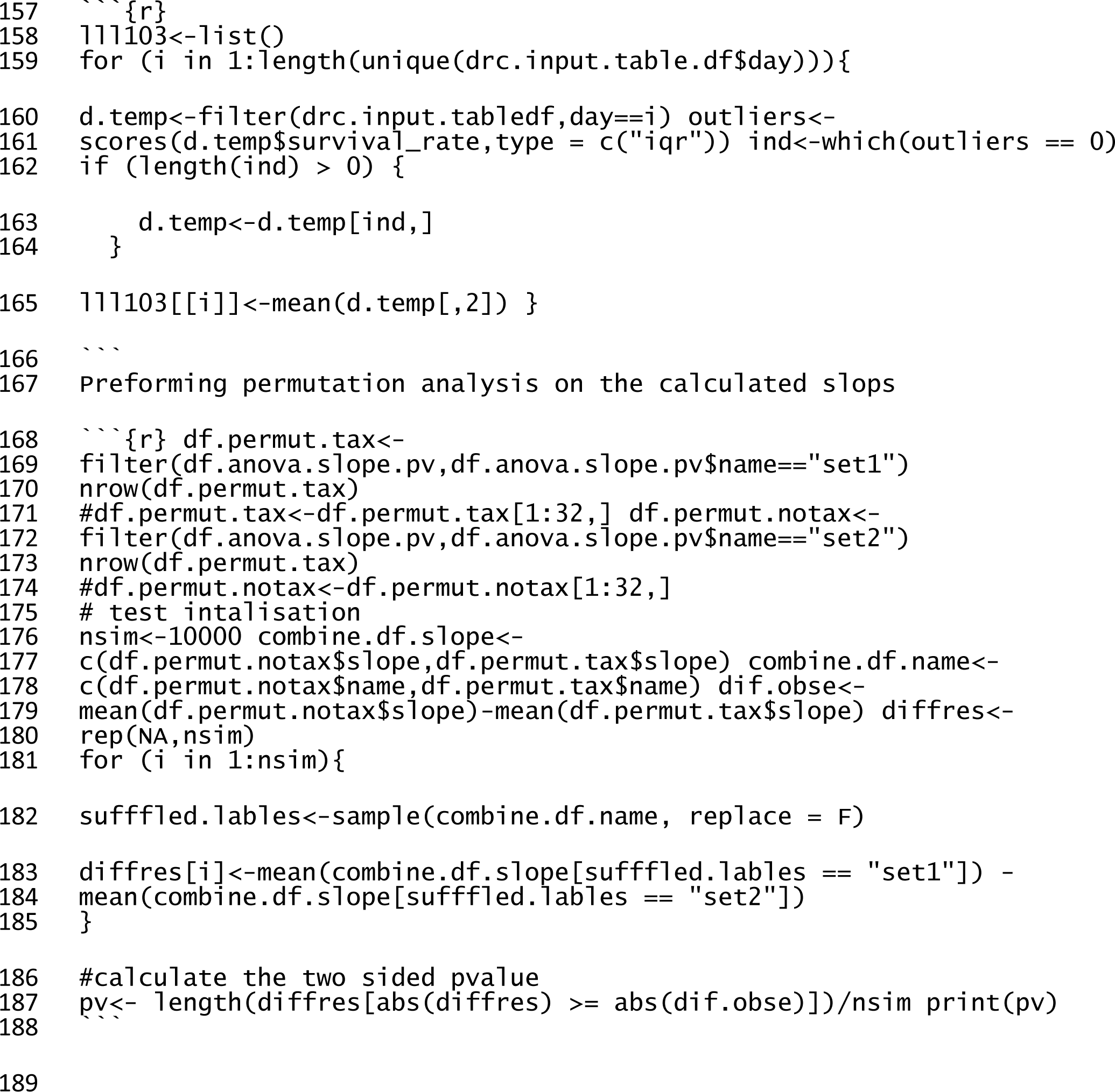

**Supplementary Table S1.**
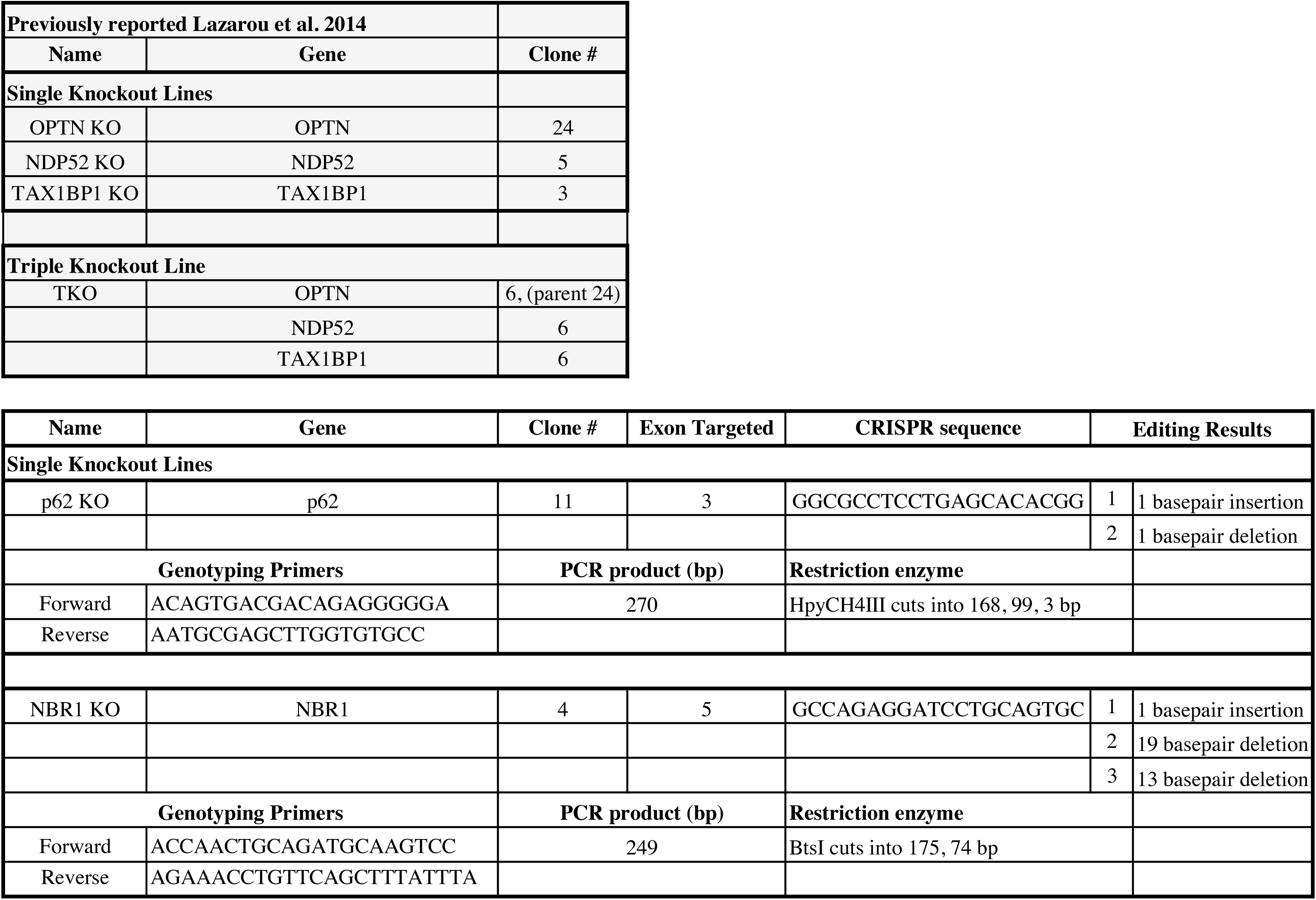

